# Programmable Cleavage of Double-stranded DNA by Combined Action of Argonaute CbAgo from *Clostridium butyricum* and Nuclease Deficient RecBC Helicase from *E.coli*

**DOI:** 10.1101/2021.07.01.450725

**Authors:** Rita Vaiskunaite, Jogirdas Vainauskas, Janna Morris, Vladimir Potapov, Jurate Bitinaite

## Abstract

Prokaryotic Argonautes (pAgos) use small nucleic acids as specificity guides to cleave single-stranded DNA at complementary sequences. DNA targeting function of pAgos creates attractive opportunities for DNA manipulations that require programmable DNA cleavage. Discovery of mesophilic Argonautes active at physiological temperature places pAgos closer to their possible application for genome editing as a simpler alternative to CRISPR/Cas nucleases. Currently, the use of mesophilic pAgos as programmable DNA endonucleases is hampered by their poor action on double-stranded DNA (dsDNA), mainly due to their inability to invade the DNA duplex. The present study demonstrates that efficient *in vitro* cleavage of double-stranded DNA by mesophilic Argonaute CbAgo from *Clostridium butyricum* can be activated via the DNA strand unwinding activity of nuclease deficient mutant of RecBC DNA helicase from *Escherichia coli* (referred to as RecB^exo-^C). Properties of CbAgo and characteristics of simultaneous cleavage of complementary DNA strands in concurrence with DNA strand unwinding by RecB^exo-^C were thoroughly explored using 0.3-25 kb DNA substrates. When combined with RecB^exo-^C helicase, CbAgo was capable of cleaving target sequences located 11-12.5 kb from the ends of linear dsDNA at 37ºC. Our study demonstrates that CbAgo with RecB^exo-^C can be programmed to generate dsDNA fragments flanked with custom-designed single-stranded overhangs suitable for ligation with compatible DNA fragments. At present, the combination of CbAgo and RecB^exo-^C represents the most efficient mesophilic DNA-guided DNA-cleaving programmable endonuclease for use in diagnostic and synthetic biology methods that require sequence-specific nicking/cleavage of dsDNA at any desired location.

## INTRODUCTION

Argonaute proteins have the ability to bind small single-stranded 5’-phosphorylated nucleic acids which provide base-pairing specificity for cleavage of complementary single-stranded targets. Eukaryotic Argonautes (eAgo) are essential components of the RNA-induced gene silencing processes (1, 2). The mechanism involves the association of eAgo with a single-stranded RNA guide to form an RNA-induced silencing complex (RISC) (3, 4). The RISC is directed to the complementary sequence on mRNA where the Argonaute cleaves single-stranded mRNA in a guide specific manner resulting in a reduction of target gene expression (5–7). RISC also can interact with a variety of Argonaute-associated proteins to induce cleavage-independent mechanisms of gene regulation (8, 9). Prokaryotes lack RNA interference pathways; however, many bacteria and archaea possess Argonaute proteins implying a different biological role for these proteins. Multiple recent studies suggest that prokaryotic Argonautes (pAgos) function *in vivo* as defense systems against foreign genetic elements (10–14). pAgos represent a very diverse group of proteins and the presence of four common domains (N, PAZ, MID and PIWI) divides them into two major groups: the short pAgos and the long pAgos (10, 12, 15). Short pAgos lack the N and PAZ domains. Long pAgos, similar to eAgos, contain all four common domains and encompass all known active pAgos (12). In contrast to eAgos, which exclusively use RNA guides to exclusively target RNA, bacterial Agos have been shown to bind either RNA or DNA guides and to cleave either RNA or DNA targets (16–19). Some archaeal Agos exclusively utilize DNA guides for cleavage of DNA targets (20, 21). Many pAgos exhibit a non-specific nuclease activity when they are not associated with guides (22–24), which has been implicated in cellular function required for guide processing (22). TtAgo co-purifies with DNA sequences that are preferentially derived from its own expression plasmid, but only if the Argonaute is catalytically active (25). The recent study of mesophilic bacterial Argonaute CbAgo from *Clostridium butyricum* shows that CbAgo nucleolytic activity cooperates with cellular double-strand break repair machinery in the generation of small DNAs that can later be used as guides by this Argonaute (14).

The ability to direct DNA guides for cleavage of complementary DNA target opens an opportunity for using pAgos as programmable DNA endonucleases (26). So far, the CRISPR/Cas9 system is the most widely used enzymatic tool for programmable DNA cleavage. Cas9 nuclease programmed with RNA guide can invade double-stranded DNA (dsDNA) and generate double-strand breaks at a guide-specific location. The CRISPR/Cas9 systems function at physiological temperatures, so they have been successfully adapted for genome editing *in vivo* (reviewed in 27). In contrast to CRISPR-Cas9, the cleavage of double-stranded targets by pAgos can proceed only if DNA strands are separated beforehand. Argonaute from the archeon *Pyrococcus furiosus* (PfAgo) was shown to work as a programmable DNA-guided DNA-cleaving endonuclease *in vitro*, but only because PfAgo is active at a temperature (>87ºC) that causes thermal DNA denaturation (28). After DNA melting takes place, the double-strand cleavage by PfAgo still proceeds by way of two independent strand-nicking events catalyzed by two PfAgo monomers loaded with separate guides that are complementary to the opposing DNA strands (28). In fact, all characterized pAgos that function at lower temperature range (30-75ºC) show low levels of endonucleolytic activity on dsDNA and preferentially cleave negatively supercoiled plasmids and/or DNA sections with low G/C content (24, 25, 29-32) consistent with the greater single-stranded character of these substrates.

Bioinformatics analysis of pAgo operons reveals the abundance of genes encoding diverse range of putative nucleases, helicases and DNA-binding proteins (10, 15, 33). Their possible connection remains unclear, but if pAgos are involved in chopping foreign dsDNA then they very likely collaborate with other cellular proteins that can open the DNA duplex. Such theoretical speculation gained some experimental support in the case of thermophilic Argonaute from *Thermus thermophilus* (TtAgo) which showed an elevated activity on dsDNA when cleavage was carried out in the presence of single-strand binding protein ET SSB or TthUvrD DNA helicase (29). Recently, dsDNA cleavage by the mesophilic Argonaute from cyanobacterium *Synechococcus elongatus* (SeAgo) was tested during ongoing transcription of a target region which was expected to transiently melt dsDNA, but the approach had no effect on target cleavage (32).

DNA helicases represent a large and diverse group of proteins that are involved in variety of biologically important cellular processes (34). The majority of DNA helicases require either a 5’- or 3’-single-stranded DNA (ssDNA) end as an initiation point to start unwinding DNA duplex and typically do not bind to blunt-ended dsDNA. One exception is *Escherichia coli* RecBCD DNA helicase which prefers unwinding blunt-ended DNA and is inhibited by ssDNA ends that exceed ~25 nucleotides (35). The RecBCD enzyme remains the fastest (1,000 to 2,000 bp s^−1^) and most processive (~30,000 bp) helicase reported in the literature (36). Wild type RecBCD is a heterotrimer consisting of three subunits, RecB, RecC and RecD and has multiple enzymatic activities: ATP-dependent DNA unwinding activity, ATP-dependent dsDNA and ssDNA exonuclease activity, ATP-stimulated ssDNA exonuclease activity and ATPase activity (36, 37). The RecB subunit is organized into a 100-kDa N-terminal helicase domain and 30-kDa C-terminal exonuclease/endonuclease domain (38). The C-terminal domain of RecB subunit is responsible for all nuclease activities associated with the RecBCD complex (39, 40). Crystal structure of RecBCD enzyme revels four conserved catalytic residues (Glu1020, Asp1067, Asp1080 and Lys 1082) in the active site of RecB nuclease domain (41). Nuclease activity of RecBCD enzyme can be completely inactivated by mutation of either aspartate residue to alanine or lysine residue to glutamine (40).

In the present study we report that mesophilic Argonaute can cleave dsDNA during concurrent unwinding of DNA strands by the nuclease deficient mutant of RecBC DNA helicase referred here as RecB^exo-^C. In total, we screened 20 putative pAgos for cleavage activity at 37-65ºC and found 10 candidates that were active at 37ºC. Six mesophilic Argonautes originating from *Clostridia* class bacteria were further biochemically characterized. While our studies were underway several other reports were published describing characterization of mesophilic Argonautes, including CbAgo from *Clostridium butyricum*, CpAgo from *Clostridium perfringens* and IbAgo from *Intestinibacter bartlettii* (24, 30, 31, 42). In our study, the most active Argonaute, CbAgo was paired with RecB^exo-^C helicase to synchronize the DNA duplex unwinding with guided cleavage of individual DNA strands at 37ºC. A detailed analysis of dsDNA cleavage by CbAgo was carried out using high-throughput capillary gel electrophoresis (CE) (29, 43). Herein, we show that in the presence of RecB^exo-^C helicase CbAgo efficiently cleaves linear dsDNA ranging from 300 bp to 25 kb in length and produces DNA fragments flanked with unique single-stranded overhangs that can be ligated to other DNA fragments with complementary overhangs. Currently, a combination of CbAgo and RecB^exo-^C represents the most efficient DNA-guided DNA-cleaving programmable endonuclease which can be used for the development of new diagnostic and synthetic biology tools that require sequence-specific nicking/cleavage of dsDNA at otherwise inaccessible locations at mesophilic temperatures.

## RESULTS

### The search for highly active mesophilic DNA-guided DNA-cleaving Argonautes

From the compiled list of 45 bacterial Agos, we selected 20 candidates to screen for catalytically active proteins (Supplementary Table S1). A quick examination of cleavage activity was performed with candidate pAgo proteins expressed with PURExpress In Vitro Protein Synthesis kit. SDS-PAGE analysis confirmed detectable levels of soluble proteins for 19 pAgo candidates, except for CbAgo protein which was predominantly found in an insoluble fraction. Under current conditions, ten pAgo candidates displayed DNA cleavage activity at 37°C and 13 candidates were active at 65°C (Supplementary Figure S1). Four pAgo candidates with the highest cleavage activity were found in hosts that belong to the *Clostridium* genus. CbAgo displayed low activity at 37°C, which was attributed to the low level of soluble protein in PURExpress sample. Interestingly, CbAgo protein was found in a soluble fraction when expressed *in vivo* in T7 Express LysY/I^q^ *E*.*coli* (Supplementary Methods).

We purified and characterized six pAgos with a goal to identify candidate with the highest activity at 37°C. Five selected pAgos originated from hosts delegated to *Clostridium* genus. The host for IbAgo originally was assigned as *Clostridium bartlettii*, but later the species was re-assigned as *Intestinibacter bartlettii* (44). We used a previously described high-throughput CE-based assay (29, 43) to rapidly characterize the purified pAgos for guide preference (DNA vs RNA) and target preference (DNA vs RNA). The results are summarized in Supplementary Table S2. All candidate pAgos preferred DNA guides and DNA targets over RNA guides and RNA targets. CpAgo was the only Argonaute capable of DNA-guided cleavage of RNA and RNA-guided cleavage of DNA, albeit at a reduced rate if compared to the DNA-guided DNA cleavage. Candidate pAgos were capable of cleaving DNA at temperatures spanning from 30°C to 75°C. The highest activity was observed at 55-65°C, except for IbAgo which was the most active at 45°C. The efficiency of DNA target cleavage at 37°C was evaluated at different time points ranging from 5 to 120 minutes. The results presented in Figure 1 demonstrate that under current conditions the candidate pAgos can be ranked in the following order: CbAgo > CpAgo > CaAgo > CdAgo > IbAgo > CsAgo. Because of the robust activity CbAgo was further investigated for ability to cleave double-stranded DNA cleavage at 37°C.

**Figure 1.**
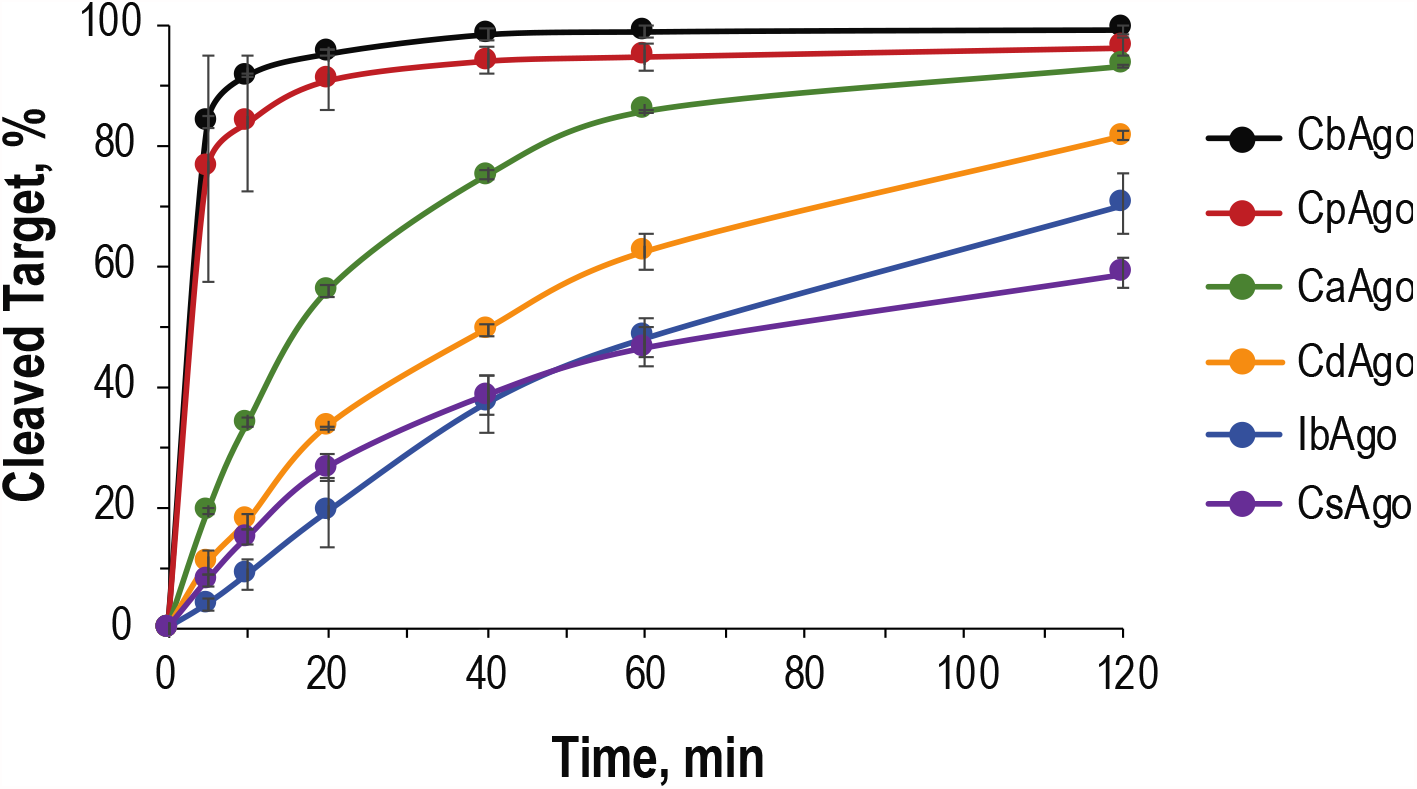
Comparison of *Clostridia* Argonautes for cleavage efficiency of ssDNA target. Cleavage reactions were performed at a final 125:125:50 nM molar concentration ratio of Ago:Guide:Target as described in Materials and Methods. Error bars indicate the standard deviation of three independent experiments. Bacterial hosts of pAgo proteins used in this study: *C*.*butyricum* (CbAgo), *C*.*perfringens* (CpAgo), *C*.*saudiense* (CaAgo), *C*.*disporicum* (CdAgo), *I*.*bartlettii* (IbAgo), *C*.*sartagoforme* (CsAgo).

### RecB^exo-^C DNA helicases assist CbAgo in cleaving targets on dsDNA

We have constructed a nuclease deficient variant of RecB helicase, referred to as RecB^exo-^, by replacing three catalytic residues, E1020, D1080 and K1082 with alanine residue. It has been reported that RecB subunit alone is a weak DNA helicase, but it interacts with RecC protein to form a rapid and processive DNA helicase (37). Therefore, individually purified RecB^exo-^ and RecC subunits were mixed at 1:1 stoichiometry to reconstitute RecB^exo-^C helicase. DNA unwinding activity of either RecB^exo-^ or RecB^exo-^C was tested using the CE-based assay as described in Supplementary Methods. The obtained results confirmed that RecB^exo-^ alone is a weak DNA helicase, but its unwinding activity significantly increases after interaction with RecC protein (Supplementary Figure S2). Nuclease deficient RecB^exo-^C helicase was then explored as a CbAgo partner in cleavage of dsDNA at physiological temperature.

To investigate CbAgo cleavage activity on dsDNA, the 21-nt long guides T2 and B2 were designed to target opposing DNA strands in the middle of a 322 bp 5’-FAM/ROX labeled DNA substrate (Figure 2A). The targeted DNA region contained a BbvCI restriction site allowing use of BbvCI restriction fragments as internal markers to verify the size of CbAgo cleavage products. In the absence of RecB^exo-^C helicase no cleavage was observed after DNA was treated with either CbAgo/T2 or CbAgo/B2 (Figure 2B, Lanes 2 and 4, respectively). The result was expected as CbAgo by itself cannot initiate guide-specific cleavage of duplex DNA. However, when CbAgo reaction was supplemented with RecB^exo-^C, either CbAgo/T2 or CbAgo/B2 complex cleaved 322 bp DNA in a guide-specific mode as confirmed by the appearance of 155 nt 5’-FAM labeled and 167 nt 5’-ROX labeled cleavage products (Figure 2B, Lanes 3 and 5). The size of the CbAgo generated fragments was confirmed by the BbvCI cleavage which produced 155 nt and 164 nt restriction fragments, respectively (Figure 2B, Lane 6). An array of 21-nt long DNA guides was used to further evaluate CbAgo efficiency for cleavage of single strands within 322 bp dsDNA in the presence of RecB^exo-^C helicase. Four guides designed to cleave a 5’-ROX labeled strand (Supplementary Figure S3A) resulted in very efficient CbAgo/guide complexes as 50-80% of the respective targets were cleaved in the first four minutes and 95-100% cleavage was achieved after 16 minutes at 37°C (Supplementary Figure S3B). Thirteen guides were designed to hybridize with the 5’FAM-labeled strand at sequences that were shifted by one nucleotide with respect to each other (Supplementary Figure S4A). This strand was cleaved significantly slower and only nine CbAgo/guide complexes could cleave the respective targets more than 50% in 16 minutes (Supplementary Figure S4B). Except for the guide T1-C1, CbAgo loaded with different guides cleaved the respective targets up to 80-100% in 64 minutes (Supplementary Figures S4). Despite the observed differences, the results provided a strong evidence that CbAgo can rapidly find and cleave transiently formed ssDNA targets imbedded on long dsDNA during ongoing strand unwinding by RecB^exo-^C helicase.

**Figure 2.**
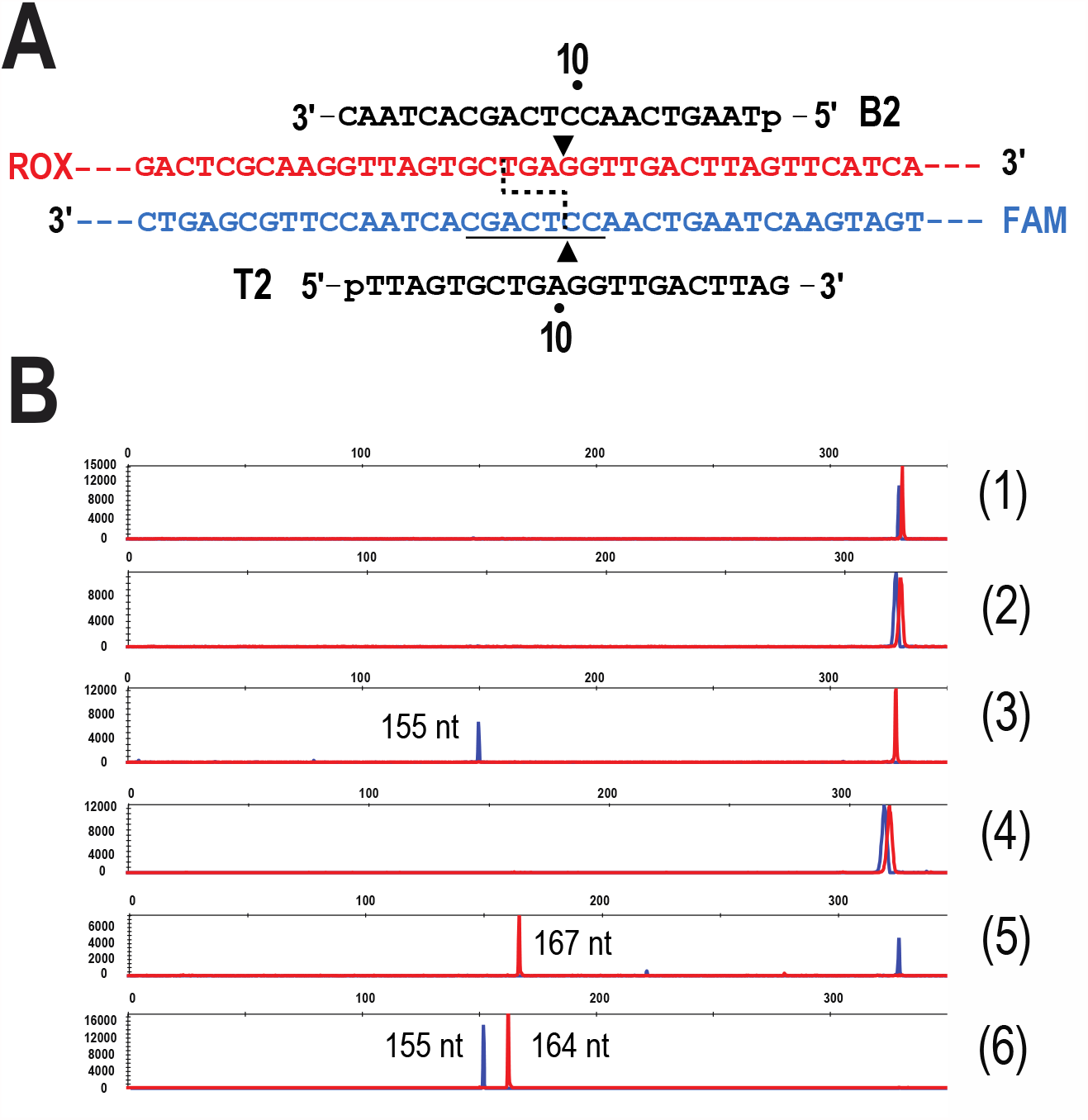
Guide-specific cleavage of double-stranded DNA by CbAgo in the presence of RecB^exo-^C helicase. **A)** Schematic overview of guide positioning on 5’-FAM/ROX labeled 322 bp dsDNA substrate. Black triangles indicate cleavage positions of CbAgo/B2 and CbAgo/T2 complexes. BbvCI recognition site is underlined, and dashed line shows BbvCI cleavage positions. (**B**) Capillary electrophoresis results. **Lane 1**, dsDNA substrate. **Lane 2**, dsDNA cleaved with CbAgo/T2. **Lane 3**, dsDNA cleaved with CbAgo/T2 in the presence of RecB^exo-^C. **Lane 4**, dsDNA cleaved with CbAgo/B2. **Lane 5**, dsDNA cleaved with CbAgo/B2 in the presence of RecB^exo-^C. **Lane 6**, dsDNA cleaved with 10 units of BbvCI restriction endonuclease.

### Effect of CbAgo:guide molar ratio on double-strand DNA cleavage

Two guide pairs were initially selected to test double-strand cleavage of a 322 bp dsDNA in the presence of RecB^exo-^C helicase. Double-strand cleavage by CbAgo loaded with guides T2 and B1 was expected to yield DNA fragments tailed with 5nt-long 3’-ss overhangs, whereas CbAgo loaded with guides T1-C2 and B4 was expected to yield DNA fragments tailed with 5nt-long 5’-ss overhangs (Left panels in Figure 3). CbAgo was pre-loaded with individual guides, then two CbAgo/guide complexes were combined in the reaction with dsDNA and RecB^exo-^C helicase and cleavage of each DNA strand was monitored over time. Concurrent cleavage of DNA strands by CbAgo/T2 and CbAgo/B1 rapidly advanced to completion (Figure 3A). Unexpectedly, a ~10-fold decline in cleavage of either DNA strand was observed when CbAgo was loaded with T1-C2 and B4 guides which were arranged to yield a 5’-staggered double-strand break (Figure 3B). The cleavage of either DNA strand remained profoundly incomplete after 64 minutes in contrast to the respective single-guided cleavage reactions (Supplementary Figures S3 and S4). To further explore the observed bias, CbAgo efficiency was compared in a series of experiments using two sets of guide pairs created by combining guides B1 and B4 targeting a 5’-ROX labeled strand with thirteen guides targeting the 5’-FAM labeled DNA strand. The use of 13 guides in combination with the guide B1 allowed to generate CbAgo cleavage products tailed with 3’-ss overhangs varying from 1 to 13 nt in length. When thirteen guide pairs were formed using guide B4, CbAgo cleavage was expected to yield products tailed with 5’-ss overhangs varying from 2 to 14 nt in length (Figure 4A). CbAgo efficiently cleaved dsDNA when loaded with guides programmed to create 3’-staggered double-stranded breaks resulting in 80-100% DNA strand cleavage after 16 minutes at 37°C (Figure 4B). However, strand cleavage was greatly reduced when CbAgo was loaded with guides arranged to generate cleavage products with 5’-ss overhangs (Figure 4C). The largest decline was observed with guide pairs positioned to yield 5’-ss overhangs varying from 5 nt to 8 nt in length, although CbAgo activity partially recovered when a 5’-ss overhang was increased from 9 nt up to 14 nt (Figure 4C).

**Figure 3.**
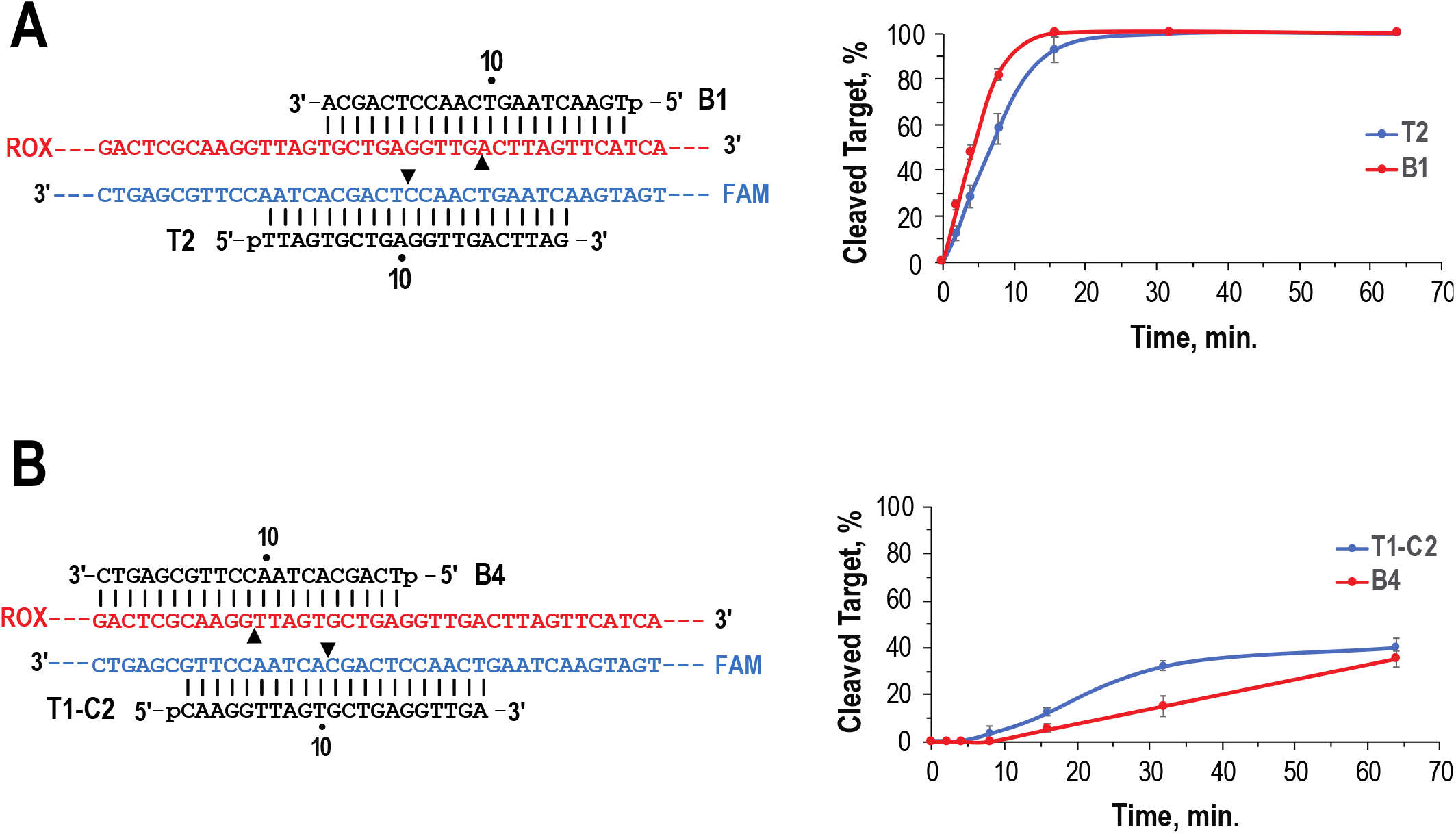
Double-strand cleavage of 5’-FAM/ROX labeled 322 bp DNA by CbAgo loaded with guide pair T2+B1 (A) or T1-C1+B4 (B). **P**anels on the left show targeted region of 322bp dsDNA substrate and guides used for cleavage of DNA strands. Black triangles show target cleavage sites. dsDNA cleavage was carried out at 37ºC and samples were removed from the reaction after 2, 4, 8, 16, 32 and 64 minutes. Cleavage products were analyzed by CE and quantified as described in Materials and Methods. Error bars indicate standard deviation of three independent experiments. **(A)** Time course of double-strand cleavage with CbAgo/T2+B1 which generates DNA fragments flanked with 5-nt long 3’-ss overhangs. (**B**) Time course of double-strand cleavage with CbAgo/T1-C2+B4 which generates DNA fragments flanked with 5-nt long 5’-ss overhangs.

**Figure 4.**
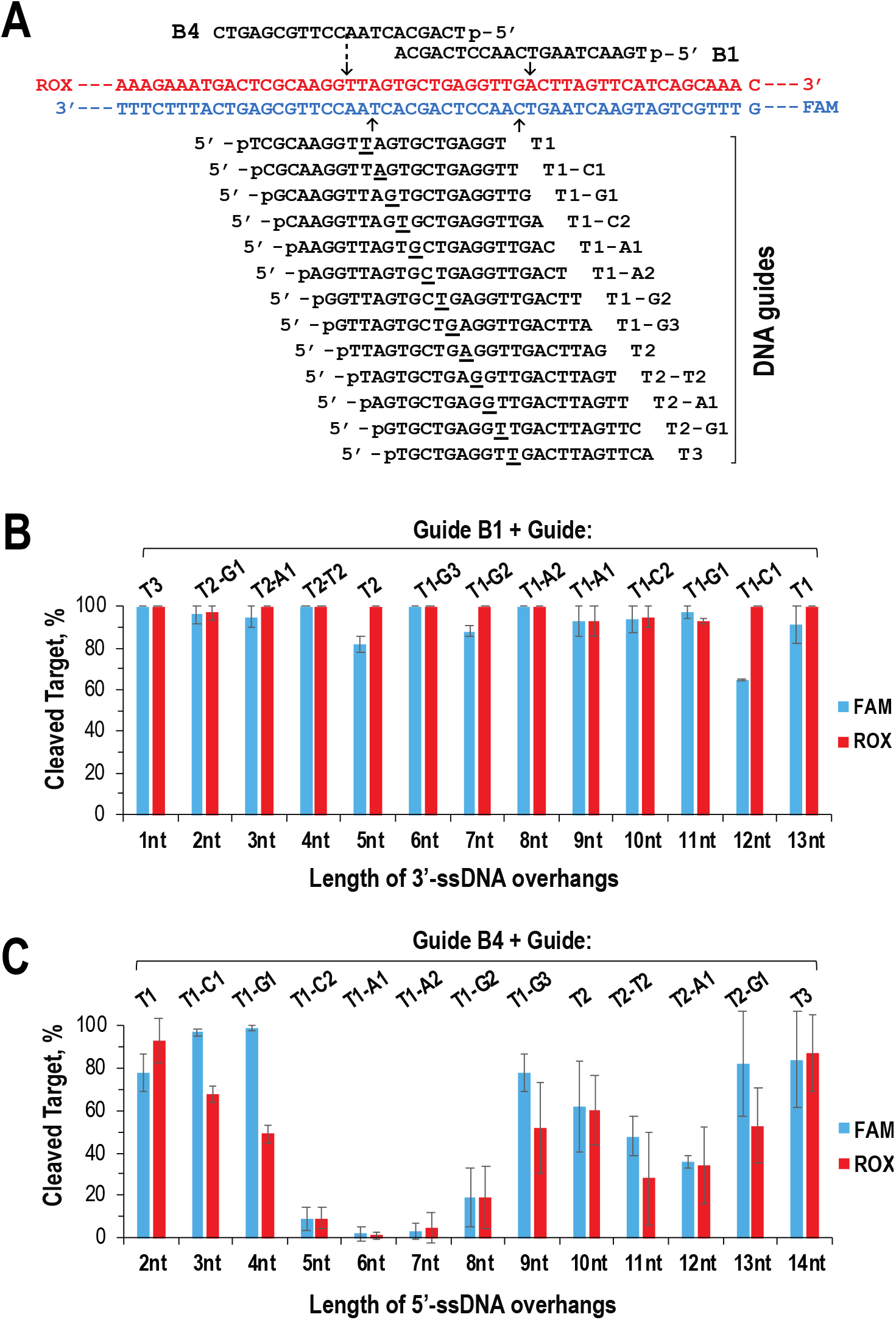
Double-strand cleavage of 5’-FAM/ROX labeled 322 bp DNA by CbAgo loaded with two sets of 13 guide pairs which generate cleavage products tailed with either 3’- or 5’-ss overhangs. **(A)** Schematic overview of guide positioning on DNA target. On the 5’-FAM labeled strand, arrows indicate cleavage position of the first (CbAgo/T1) and the last (CbAgo/T3) complexes. CbAgo/guide complexes were combined with dsDNA substrate at CbAgo:guide:target molar ratio of 125:250:50 nM. The concurrent double-strand cleavage was carried out for 16 minutes at 37ºC in the presence of 250 nM RecB^exo-^C. The percentage of cleaved DNA was quantified for each DNA strand. **(B)** Efficiency of double-strand cleavage by CbAgo loaded with the indicated guide pairs which generate cleavage products tailed with 3’-ss overhangs varying from 1 to 13 nt in length. Error bars indicate the standard deviation of two independent experiments. **(C)** Efficiency of double-strand cleavage by CbAgo loaded with the indicated guide pairs which generate cleavage products tailed with 5’-ss overhangs varying from 2 to 14 nt in length. Error bars indicate the standard deviation of three independent experiments.

Under current reaction conditions, CbAgo was loaded with each guide at a 1:2 molar concentration ratio with intent to evade a non-specific DNA “chopping” by a guide-free CbAgo (23–25). This indicates that double-strand cleavage is carried out in the presence of free guides that are highly complementary to each other. We then postulated that free guides potentially may interfere with dsDNA cleavage when guides are arranged to yield 5’-staggered double-stranded breaks. To verify this prediction, a detailed analysis of varying CbAgo:guide molar concentration ratios was performed using a T1-A1+B4 guide pair (Figure 5). Double-strand cleavage by CbAgo/T1-A1+B4 was nearly eliminated when CbAgo was loaded with guides at molar ratios ranging from 1:2 (125:250 nM) to 1:1.6 (125:200 nM). Interestingly, CbAgo activity partially recovered when Ago:guide molar ratio was reduced to 1:1.4 (125:175 nM), and at a 1:1 molar ratio (125:125 nM) more than 90% cleavage of both strands was achieved after 16 minutes at 37°C (Figure 5). Next, CbAgo was loaded at the 1:1 molar ratio with 13 guide pairs arranged to produce ds fragments with 5’-ss overhangs of varying length. Remarkably, an efficient cleavage of both DNA strands was observed with all guide pairs when cleavage reactions were performed using equimolar CbAgo and guide concentrations (Supplementary Figure S5). The results clearly indicated that CbAgo activity was inhibited by the presence of free guides in the reaction, but only if the guides were arranged to yield products with 5’-ss overhangs. In contrast, no inhibition occurred if a 2-fold molar excess of guides over CbAgo was used to generate double-stranded cleavage products tailed with 3’-ss overhangs (Figure 4B). In summary, we demonstrated here that in the presence of RecB^exo-^C helicase CbAgo can efficiently cleave dsDNA and produce DNA fragments flanked with either 3’- or 5’-ss overhangs of varying length, but a proper Ago:guide molar ratio must be used when aiming to create cleavage products tailed with 5’-ss overhangs.

**Figure 5.**
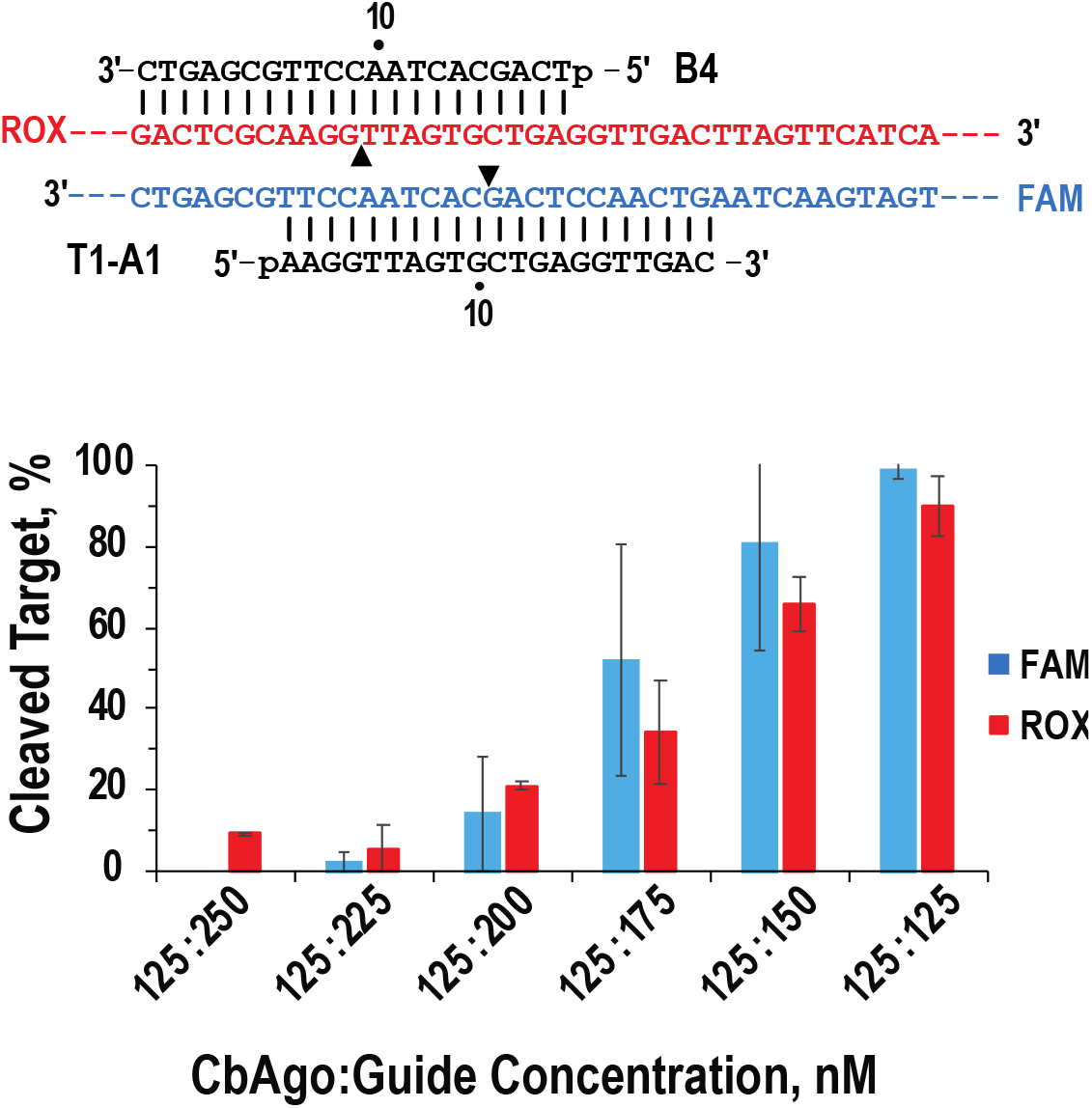
Efficiency of double-strand cleavage of 5’-FAM/ROX labeled 322 bp DNA at different CbAgo:guide molar concentration ratios. CbAgo/T1-A1 and CbAgo/B4 complexes were combined with DNA target at CbAgo:guide:target molar concentration ratios of 125:250:50, 125:225:50, 125:200:50, 125:175:50, 125:150:50 and 125:125:50 nM. Double-strand cleavage was carried out for 16 minutes at 37ºC in the presence of 250 nM RecB^exo-^C helicase and the percentage of cleaved DNA was quantified for each DNA strand. Error bars indicate standard deviation of three independent experiments.

### CbAgo acts as programmable DNA-guided endonuclease on linear double-stranded DNA

The linearized 5.4 kb φX174 phage DNA was used to explore if CbAgo/RecB^exo-^C can cleave targets on substantially longer dsDNA. Four targets were selected for cleavage, where each target was positioned in the middle of the properly linearized φX174 DNA, therefore in all cases CbAgo was expected to generate two identical size DNA fragments that could be detected by agarose gel-electrophoresis (Figure 6A). Results demonstrated that in all four cases the opposing CbAgo/guide complexes were capable of binding and cleaving transiently formed single-stranded targets on both DNA strands once RecB^exo-^C unwound the 5.4 kb DNA (Figure 6A, lane 2 in panels I-IV). The specificity of DNA-guided cleavage was confirmed by cleaving a SspI-linearized φX174 DNA with SapI which produced restriction fragments similar in size to fragments generated by CbAgo/T2+B1 (Figure 6A, lane 3). DNA cleavage by CbAgo programmed with either SapT+SapB or StuT+StuB guide pairs (panels III and IV) was less efficient than at the other two locations that were targeted with guide pairs T2+B1 and XhoT+XhoB (panels I and II). All guide pairs were designed to generate 5nt-long 3’-ss overhangs at the cut site and all guide sequences started with a 5’-pT terminal nucleotide. In single-guided reactions, the CbAgo cleavage efficiency was found to be dependent on a guide sequence (Supplementary Figure S4). Likewise, the preference for guides with a certain nucleotide sequence could be behind the observed differences in CbAgo cleavage of φX174 DNA, thus implying that optimization of guide sequences might be necessary to achieve a complete target cleavage at selected locations.

**Figure 6.**
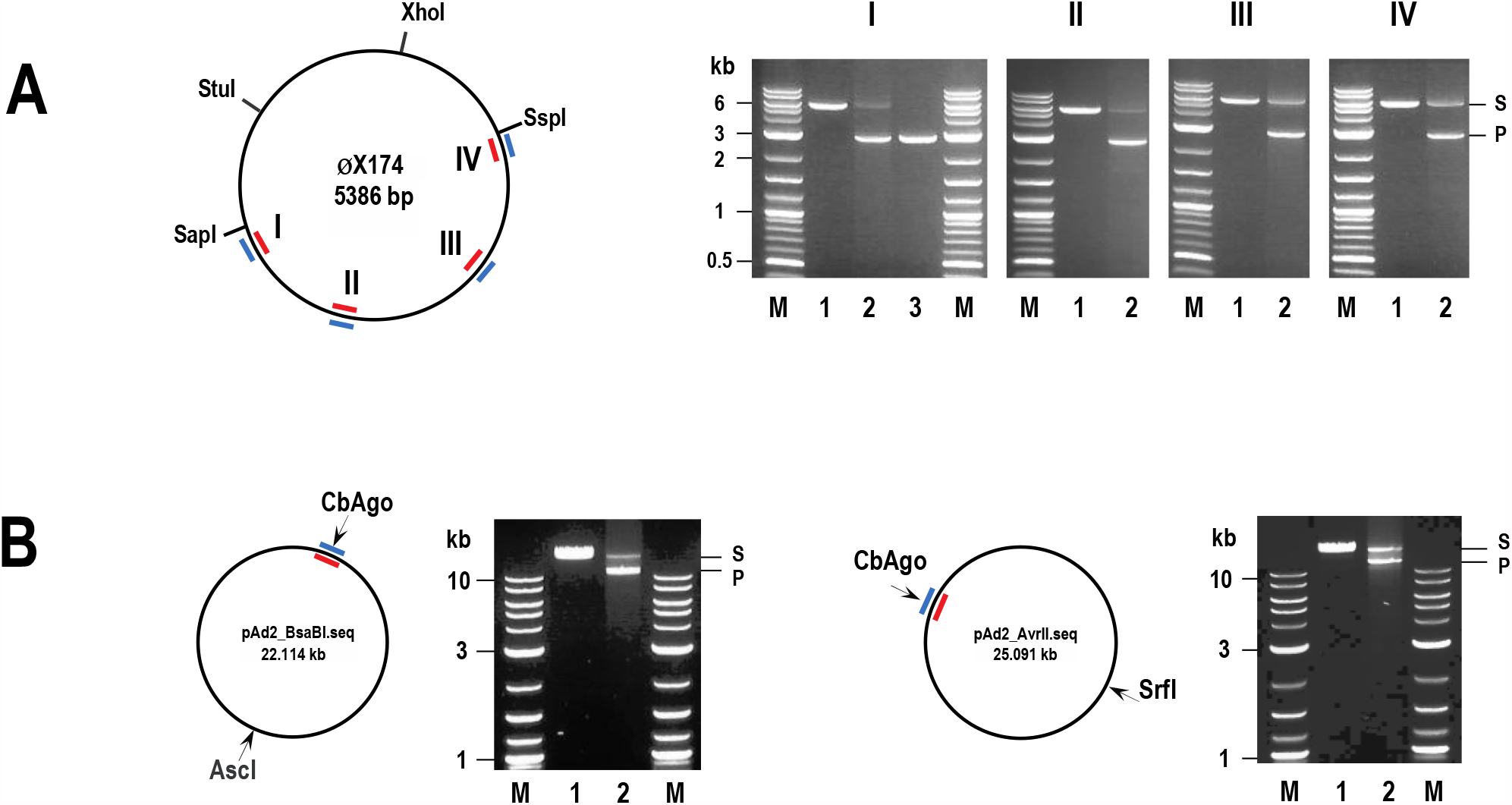
CbAgo acts as a programmable DNA endonuclease on linear dsDNA. The CbAgo cleavage was carried out either in the absence (Lane 1) or in the presence of RecB^exo-^C (Lane 2). The circular map of each plasmid with restriction sites and guide positions is shown on the left of the respective gel. (**A**) Four locations of linear φX174 DNA were targeted with CbAgo loaded with four different guide pairs. **Panel I**, SspI-linearized DNA cleaved with CbAgo/T2 and CbAgo/B1. Lane 3, SspI+SapI digested DNA. **Panel II**, XhoI-linearized DNA cleaved with CbAgo/XhoT and CbAgo/XhoB. **Panel III**, StuI-linearized DNA cleaved with CbAgo/StuT and CbAgo/StuB. **Panel IV**, SapI-linearized DNA cleaved with CbAgo/SapT and CbAgo/SapB. (**B**) AscI-linear pAd2_BsaBI DNA (22.1 kb) cleaved with CbAgo/AdB-450T and CbAgo/AdB450B. (**C**) SrfI-linearized pAd2_AvrII DNA (25.1 kb) cleaved with CbAgo/AdA-20860T and CbAgo/AdA-20860B. S, linear DNA; P, CbAgo cleavage products; M, 1kb plus DNA ladder.

A mesophilic Ago activity has never been demonstrated on dsDNA longer than 5-6 kb. To determine if CbAgo/ RecB^exo-^C combination was able to cleave 20-25 kb DNA at 37ºC, the plasmids pAd2_BsaBI (22.114 kb) and pAd2_AvrII (25.091 kb) were linearized with AscI and SrfI, respectively, and two guide pairs were designed to target each DNA at a midpoint (Figure 6B). In the presence of RecB^exo-^C, CbAgo loaded with the respective guide pair specifically cleaved targeted sites as indicated by the appearance of 11 kb and 12.5 kb cleavage products (Figure 6B, lane 2). Remarkably, the concurrent DNA unwinding by RecB^exo-^C permits CbAgo to cleave DNA targets located at 11-12.5 kb distance from the end of dsDNA implying that CbAgo/guide complex can find and bind transiently formed single-stranded targets faster than strand re-annealing takes place at physiological temperature.

### PCR-free method for seamless assembly of DNA fragments by using CbAgo and RecB^exo-^C

Capitalizing on the success of programmable dsDNA cleavage by CbAgo/RecB^exo-^C, we considered a method for seamless assembly of CbAgo-cleaved DNA fragments. Since Argonaute cleaves DNA strands in two independent events, a double-strand cleavage offers strategic advantages as the most efficient cleavage sites can be selected close to the end of targeted DNA and ssDNA overhangs of preferred length and composition can be created to facilitated DNA assembly. A synthetic dsDNA oligonucleotide equivalent to the two cleaved-off terminal fragments then can be used to directionally assemble two DNA fragments via complementary single-stranded overhangs (Figure 7A).

**Figure 7.**
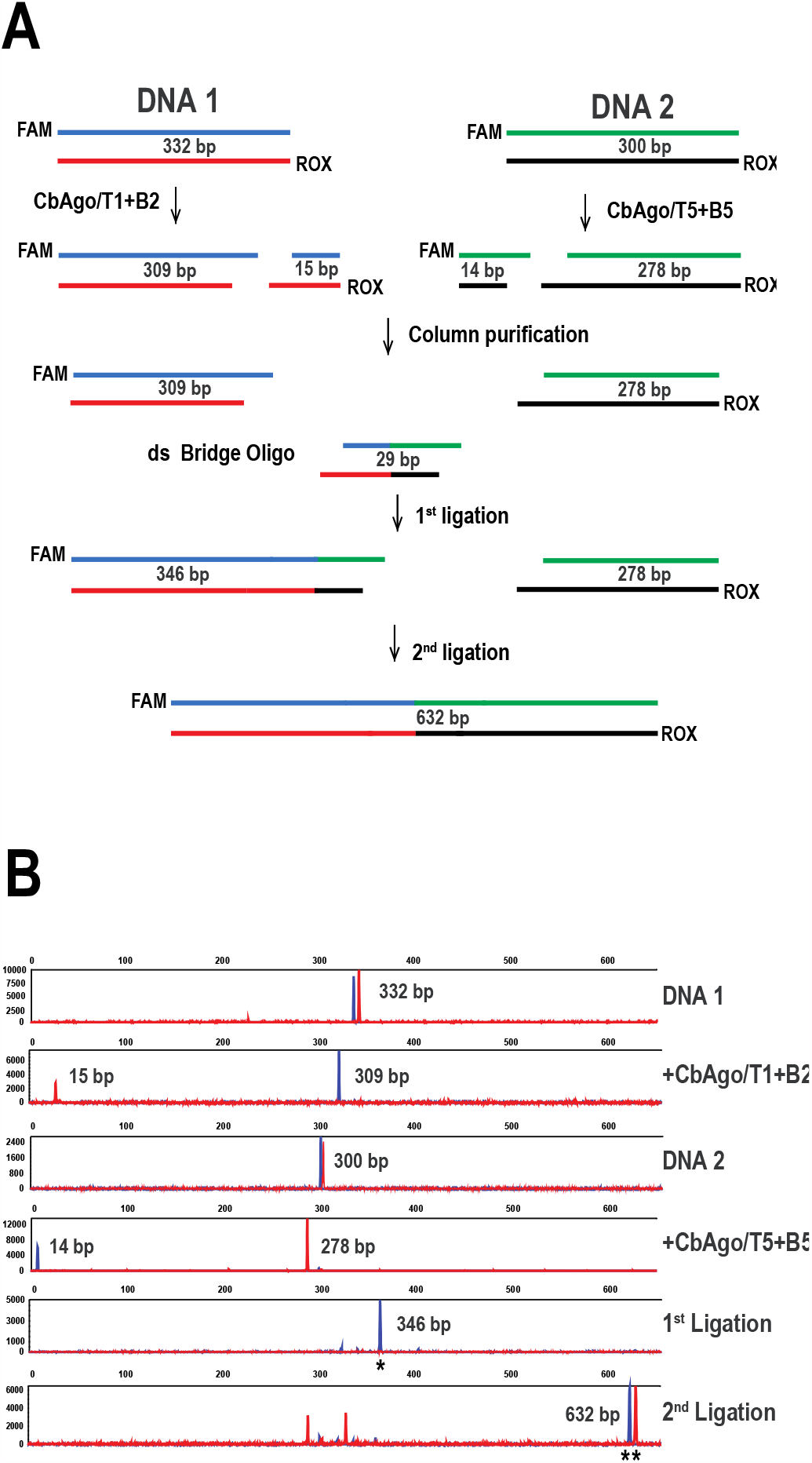
Seamless and directional assembly of dsDNA fragments using CbAgo and RecB^exo-^C. **(A)** Schematic overview of seamless DNA assembly. CbAgo/guide cleavage was carried out in the presence of RecB^exo-^C DNA helicase as described in Materials and Methods. (**B**) CE analysis of CbAgo cleavage and ligation products. Ligated fragments are marked with an asterisk (*).

As a prove-of-principal we assembled 332 bp and 300 bp 5’-FAM/ROX labeled PCR fragments (referred to as DNA1 and DNA2, respectively). On DNA1, the CbAgo target site was selected close to the 5’-ROX labeled end, whereas on DNA2 the CbAgo target site was selected close to the 5’-FAM-labeled end. The guide pairs T1+B2 and T5+B5 complemented DNA1 and DNA2, respectively, and both guide pairs were arranged to create 8nt-long 3’-ss overhangs on CbAgo cleavage products. In this arrangement, the CbAgo cleavage of either DNA produced throwaway terminal fragments, a 5’-ROX labeled 15 bp fragment and a 5’-FAM labeled 14 bp fragment, which were eliminated by column purification (Figure 7B). DNA1 and a synthetic 29 bp “Bridge” oligonucleotide were then ligated for 15 minutes at 37ºC to generate a 346 bp intermediate ligation product. The ligation reaction was then supplemented with DNA2 and ligation continued for another 15 minutes. Results presented in Figure 7B reveal the appearance of 346 bp 5’-FAM labeled ligation product during the 1^st^ ligation step, and the appearance of 632 bp 5’-FAM/ROX labeled ligation product during the 2^nd^ ligation step indicating that both CbAgo cleavage products and Bridge oligonucleotide were directionally assembled into a recombinant DNA. The results provided a compelling evidence that CbAgo cleavage generates DNA fragments with sequence-specific ssDNA overhangs that are ready for ligation without any further enzymatic treatment confirming that seamless assembly of linear DNA molecules can be accomplished using CbAgo/RecB^exo-^C programmable DNA endonuclease.

## DISCUSSION

The mechanism of double-strand DNA cleavage has mainly been reported for thermophilic pAgos which act in a range of temperatures that trigger thermal DNA denaturation. In this model, it has been shown that pAgo nuclease can cleave dsDNA in two independent events if two pAgo/guide complexes are used to target antiparallel DNA strands (28). Mesophilic pAgos act poorly on double-stranded substrates because they are unable to invade duplex DNA. Many studies have shown that only plasmids with destabilized double-stranded regions can be targeted by pAgos leading to a suggestion that *in vivo* mesophilic Agos potentially rely on natural processes that involve strand separation (24, 25, 26, 29-32). The recent study of moderately thermophilic TtAgo showed that dsDNA cleavage was stimulated by a single-strand binding protein ET SSB, and to a lesser degree by a thermophilic DNA helicase TthUvrD (29).

In this study, we explored the possibility of using a mesophilic DNA helicase for DNA unwinding to initiate individual strand cleavage at physiological temperature by mesophilic Argonaute CbAgo from *Clostridium butyricum*. Most DNA helicases require either 5’- or 3’-ssDNA end as an initiation point to start DNA duplex unwinding. At high protein to DNA concentration, EcoRecQ helicase can initiate duplex DNA unwinding from a blunt end (45, 46). Initially, we tested CbAgo in combination with EcoRecQ (Abcam, Inc., Cambridge, MA, USA) and found that CbAgo and EcoRecQ mixture was proficient in unwinding and cleaving short dsDNA. However, all efforts to cleave longer than 200 bp dsDNA substrate were unsuccessful implying that EcoRecQ is not efficient in unwinding long duplex DNA. RecBCD is the fastest and most processive DNA helicase that prefers unwinding blunt-ended DNA (36). We constructed a nuclease deficient variant of RecB helicase and reconstituted it with the RecC subunit to form a rapid and highly processive RecB^exo-^C helicase. Here we have demonstrated that DNA unwinding by RecB^exo-^C permits DNA-guided CbAgo to access targets on dsDNA substrates. The combination of RecB^exo-^C and CbAgo was successfully used to cleave linear dsDNA substrates ranging from 300 bp to 25 kb in length.

The approach of combining RecB^exo-^C with CbAgo provides an unprecedented opportunity to investigate DNA-guided cleavage of double-stranded DNA at 37ºC. Multiple CbAgo/guide complexes were compared for cleavage of either one or both DNA strands using a high-throughput CE technique. In our study, thirteen DNA guides were designed to hybridize with DNA targets located on the same strand at positions shifted by one nucleotide with respect to each other. Such guide design changed the 5’-terminal nucleotide in the guide sequence and shifted the CbAgo cleavage position by one nucleotide along the substrate DNA. The nature of the 5’-nucleotide had no significant effect on target cleavage. In addition, there was no clear evidence that cleavage efficiency was affected by the nature of nucleotides hybridizing with the target sequence in the proximity to the scissile phosphodiester bond. While the pool of tested guides was too small to credibly validate the overall effect of guide sequence on CbAgo efficiency, our results were consistent with CbAgo cleavage results previously observed on single-stranded targets (24, 31, 42). However, it is noteworthy to point out that some guides (i.e., guides T1-C1 and T2-A1) resulted in a noticeably lower cleavage efficiency if compared to adjacent guides, thus implying the efficacy of the guide might change substantially if it is moved by one nucleotide along the DNA sequence (Supplementary Figure S4). Recently a large-scale systematic study of guide preference was performed for TtAgo revealing activity correlation with the 1^st^ and 12^th^ base of the guide and to a lesser degree with the bases surrounding the cut site (47). Potentially, guide sequence preferences may be determined for CbAgo/RecB^exo-^C by performing similar large-scale studies, thus allowing to improve guide design for programmable cleavage of long dsDNA. Another interesting observation was made regarding efficiency of single-guided and double-guided cleavage of dsDNA. In single-guided reactions, CbAgo/B1 complex very rapidly cleaved the targeted 5’-ROX labeled strand (Supplementary Figure S3). In contrast, many CbAgo/guide complexes were less efficient in targeting the opposing 5’-FAM labeled strand of dsDNA (Supplementary Figure S4). However, we noticed that cleavage of the 5’-FAM labeled strand proceeded significantly faster in double-guided reactions when CbAgo/B1 was used to simultaneously cleave the opposing DNA strand (Figure 4B). Previously, ssDNA cleavage kinetics revealed product release is the rate-limiting step in the action of CbAgo (24, 30). Possibly, the slow dissociation of CbAgo/B1 complex after rapid cleavage of one DNA strand inadvertently delays re-annealing of DNA strands providing enough time for another CbAgo/guide complex to bind and cleave the opposite DNA strand.

The concurrent cleavage of DNA strands permits custom designed cleavage products choosing the length and polarity of single-stranded overhangs (i.e., 5’- or 3’-ss overhang). In our study cleavage of dsDNA was investigated using two sets of 13 guide pairs designed to generate cleavage products tailed with either 3’- or 5’-ss overhangs of varying length. CbAgo was found to efficiently cleave both DNA strands if guide pairs were prearranged to yield fragments tailed with 3’-ss overhangs, however cleavage was significantly reduced if the guide pairs were designed to generate fragments tailed with 5’-ss overhangs (Figure 4). A comprehensive study of varying CbAgo:guide molar ratios revealed that presence of free guides in the CbAgo cleavage reaction inhibited double-strand cleavage in cases when opposing CbAgo/guide complexes were arranged to yield a 5’-staggered break. Our results indicate that inhibition can be circumvented by loading CbAgo with guides at CbAgo:guide molar ratios ranging from 1:1.4 to 1:1 (Figure 5 and Supplementary Figure S5). The guides that target antiparallel DNA strands are partially complementary to each other if they are arranged to generate fragments tailed with ss overhangs shorter than 15 nucleotides (Figure 4A). Nucleotide sequence alignment of opposing guides shows that guides must have complementary 3’-terminal sequences to yield fragments with 3’-ss overhangs and complementary 5’-terminal sequences to yield fragments with 5’-ss overhangs. The 5’-terminus of a guide plays a key role in target recognition and cleavage by Ago nuclease. Nucleotides 2–8 of the guide counted from the 5’-end are termed the “seed” region. In an Ago/guide complex, the bases of the seed region are solvent exposed, therefore they can readily base pair with a matching sequence on the target strand (12, 48, 49). The crystal structure of CbAgo in complex with a guide and complementary target reveals a 15 base pair duplex formed by nucleotides 2-16 of a guide and a target (30). Furthermore, the backbone phosphates and nucleotides forming the distal 16^th^ base pair appears to be anchored by specific residues of CbAgo N-terminal domain. This implies that at least 16-nt long target is anticipated for CbAgo/guide/target complex to adopt a catalytically favored configuration. Minimal target length requirements were investigated for TtAgo loaded with a 21-nt guide DNA (16). The study showed that truncation of the target to 16 nt did not alter TtAgo cleavage activity, but 15- and 14-nt targets showed 120- and 400-fold reduced cleavage rates, respectively, and a 12-nt target was not cleaved. When two guides are arranged for a double-stranded break and provided in excess of pAgo, the seed region of one CbAgo/guide complex might base pair with the opposing free guide if the guides complement each other by 5’-terminal sequences, thus forming a “CbAgo/guide/antisense guide” complex. If 5’-complementary sequences of opposing guides are long enough, then at the initial reaction stage CbAgo/guide complexes can bind and cleave free guides instead of dsDNA targets are yet to be unwound. A delayed target cleavage of at least one DNA strand was observed with guide pairs having 5’-complementary sequences longer than 16 nucleotides. However, DNA cleavage was severely inhibited when CbAgo/guide complex and free opposing guide were capable of base pairing by 12-15 nucleotides (Figures 3B and 4C). Most likely, the guides bound as 12-15 nt long targets either were cleaved at very reduced rates or could not be cleaved at all implying that CbAgo might have similar requirements for minimal target length as TtAgo (16). Considering that product release is the limiting reaction step (24, 30), the release of the uncut 12-15 nt-long target could also be obstructed resulting in catalytically impaired “CbAgo/guide/antisense guide” complexes. For efficient double-strand cleavage CbAgo must be loaded with guides at a 1:1 molar ratio when two CbAgo/guide complexes are arranged to yield a 5’-staggered break (Figure 5). However, the use of equimolar CbAgo:guide concentration may result in unwanted appearance in the cleavage reaction of a guide-free CbAgo which may damage sequence-specific ends of dsDNA due to the inherent non-specific nuclease activity of the guide-free pAgo (22–24). This problem can be evaded by using guides programmed to yield 3’-ss staggered double-stranded breaks as efficiency of dsDNA cleavage is independent of the CbAgo:guide molar ratio when opposing guides complement each other on their 3’-ends. We have shown that CbAgo loaded with guides at a 1:2 molar ratio efficiently generates DNA fragments with sequence-specific 3’-ss overhangs ready for ligation without any further enzymatic treatment.

Two enzymatic qualities are essential for efficient cleavage of double-stranded DNA at mesophilic temperature - a rapid and processive DNA strand separation and a rapid cleavage of transiently formed ssDNA targets. We have demonstrated here that mesophilic CbAgo can efficiently cleave linear dsDNA with the help of RecB^exo-^C DNA helicase. Future research will be needed to investigate other DNA helicases and potentially other cellular proteins capable of invading duplex DNA. Ago operons reveal the abundance of genes encoding diverse range of putative nucleases, helicases, and DNA-binding proteins (10, 15, 33). Future examination of these activities potentially may lead to discovery of new cellular proteins capable of assisting pAgos in cleaving dsDNA. In our study, CbAgo was selected from six candidates as the most efficient at 37°C (Figure 1). However, under experimental conditions used to identify the optimal temperature, CbAgo was most active at 50-65°C and displayed a ~5-fold lower activity at 37°C (Supplementary Figure S6). In the past few years, the Argonaute CbAgo from *Clostridium butyricum* has become the most studied mesophilic Argonaute (14, 24, 30, 31, 42). Two highly homologous variants of CbAgo were investigated by different research groups (Supplementary Figure S7). Careful inspection of the reported temperature range revealed that another CbAgo variant displayed the highest activity at lower than 50°C temperatures and had very little or no activity at 60-64°C (30, 42). The effect of temperature on the activity of two variants was tested employing different activity assays using different guides, targets and Ago:guide:target molar ratios, all of which could significantly affect cleavage efficiency. A direct comparison of both CbAgo variants side-by-side is needed to validate if the identified differences are authentic. Nevertheless, it may be speculated that one CbAgo variant is more active at 37°C that the other as it is more evolved to function at temperature physiologically relevant to the Clostridia host. If so, then it is possible to envision that highly homologous Argonautes might display different enzymatic properties and further investigation of pAgos originating from related bacterial strains might result in finding Argonautes with properties that are highly desirable for the applied DNA manipulations.

The results presented in this study identify CbAgo/RecB^exo-^C as the most efficient mesophilic DNA-guided DNA-cleaving programmable endonuclease for *in vitro* specific nicking/cleavage of dsDNA at otherwise inaccessible locations. The most exciting conclusion of our work is that in the presence of RecB^exo-^C DNA helicase CbAgo can rapidly bind and cleave targets located as far as 11-12.5 kb away from the end of a linear dsDNA substrate. Furthermore, we have shown that CbAgo and RecB^exo-^C enzyme combination can cleave targets located close to the end of linear dsDNA and generate DNA fragments with highly specific ssDNA overhangs which are ready for ligation without further enzymatic treatment. CbAgo generated DNA fragments can be directionally and seamlessly assembled using a synthetic double-stranded oligonucleotide (Figure 7). With further optimization and improvements, a PCR-free method of seamless DNA assembly may be developed that allows to directionally join long natural DNAs. Our future studies will continue to identify accessory proteins that can aid mesophilic Agos in cleaving dsDNA *in vitro* and *in vivo*. Information gained from our current and future work might eventually lead to the development of novel tools for diagnostic and synthetic biology and possibly for genome editing that could be comparable to CRISPR-Cas mediated genome editing technologies.

## MATERIALS AND METHODS

### Bioinformatics analysis for identifying mesophilic pAgos

Candidate Argonaute proteins were identified in a series of steps. First off, three known Argonaute proteins TtAgo (UniProt ID Q746M7), NgAgo (UniProt ID L0AJX6), and PfAgo (UniProt ID Q8U3D2) were aligned using MUSCLE multiple sequence alignment software (50). The resulting multiple sequence alignment was used as an input to PSI-BLAST to search against UniProt database. The expected threshold was set at 1×10^−4^, and PSI-BLAST ran multiple iterations until convergence. All bacterial homolog hits were extracted based on taxonomic classification. Only proteins containing PAZ and PIWI domains were considered for further analysis. The presence of catalytic PIWI and PAZ domains in homologs was checked by running HMM search using domain profiles available in PFAM database (PFAM PF02171 and PF02170 for PIWI and PAZ domain, respectively). The PAZ domain profile in PFAM is built mainly on sequences of eukaryotic proteins and resulted in a very few hits when run against bacterial proteins. Therefore, the HMM profile for bacterial PAZ domain was generated from scratch using HMMER software (51) based on sequences of known Argonaute proteins. Proteins originating from known thermophilic organisms were discarded. Additionally, only proteins that share less than 90% sequence identity were selected for further analysis. Finally, proteins that did not contain aspartates at conserved PIWI catalytic sites were excluded. The remaining list of forty-five bacterial Argonautes is available in Supplementary Table S1.

### Constructs for pAgo expression and purification

The codon optimized genes encoding Argonaute proteins were ordered in pET29a expression vectors from GenScript (Piscataway, NJ, USA). Analytical amounts of twenty Argonaute proteins were synthesized from pET29a plasmids using PURExpress In Vitro Protein Synthesis kit (New England Biolabs, Inc., Ipswich, MA, USA). For large scale expression and purification of CbAgo, CpAgo, IbAgo, CdAgo, CsAgo and CaAgo, the respective genes were subcloned into pET28c expression vector in frame with the N-terminal 6xHis tag. Argonaute protein expression and purification procedures are provided in Supplementary Materials.

### Construction of RecB^exo-^ and RecC expression clones

The nuclease deficient mutant of RecB DNA helicase, referred to as RecB^exo-^, was constructed by replacing catalytic residues E1020, D1080 and K1082 with alanine residues. A plasmid encoding Exonuclease V (RecBCD) (New England Biolabs, Inc., Ipswich, MA, USA) was used as a template to amplify recB gene as three overlapping PCR fragments B1 (3080bp), B2 (200bp) and B3 (340bp). Primer sequences can be found in Supplementary Table S3. To introduce E1020A mutation, the GAG codon was replaced by GCG codon in the overlapping primers used for amplification of fragments B1 and B2. Similarly, to introduce D1080A and K1082A mutations, codon GAC was changed to GCC and codon AAA was changed to GCA in the overlapping primers for amplification of fragments B2 and B3. Three recB fragments were directly assembled into pET28c vector in frame with the N-terminal 6xHis tag employing NEBuilder HiFi DNA Assembly Cloning Kit (New England Biolabs, Inc., Ipswich, MA, USA). Wild type RecC encoding gene was individually sub-cloned into pET28c vector in frame with the N-terminal 6xHis tag. RecB^exo-^ and RecC protein expression and purification is provided in Supplementary Materials.

### Argonaute cleavage assays on single-stranded DNA or RNA substrates

All 5’-FAM labeled substrate oligonucleotides (DNA or RNA) and 5’-phosphorylated guides were purchased from Integrated DNA Technologies (Coralville, Iowa, USA). Nucleotide sequences can be found in Supplementary Table S3. To test activity of *in vitro* expressed pAgo proteins, 1 μl of PURExpress sample was mixed with 250 nM guide G-1 (21 nt) in 10 μl of buffer containing 20 mM Bis-Tris propane, pH 8.0, 50 mM NaCl, 2 mM MgCl_2_, 0.1% (v/v) Triton X-100 and incubated for 20 minutes at 37°C to form a pAgo/guide complex. The pAgo/G-1 complex was then combined with 50 nM 5’-FAM labeled target T-1 in a 20 μl reaction and incubated for 1 hour either at 37°C or at 65°C. The reactions were terminated by adding 20 μl stop buffer (95% Formamide, 0.025% Bromophenol Blue, 0.025 % Xylene Cyanol, 5mM EDTA) and heating the samples for 5 minutes at 95°C. The cleavage products were separated by gel electrophoresis on 15% denaturing polyacrylamide gel containing 7.5M urea and 24% formamide and visualized using a Typhoon 9400 Scanner (GE Healthcare Chicago, IL, USA). Activity assays performed with purified Argonaute proteins were carried out with 17 nt long guide G-2 (DNA) and G-3 (RNA). For guide loading, 125 nM Ago was combined with 125 nM guide in 95 μl of buffer containing 20 mM Bis-Tris propane, pH 8.0, 50 mM NaCl, 2 mM MgCl_2_, 0.1% (v/v) Triton X-100 and incubated for 20 minutes at 37°C. The Ago/guide mixture was combined with 50 nM 5’-FAM labeled substrate (either T-2 or T-3) in 100 μl cleavage reaction and incubated at 37°C. 10 μl samples were withdrawn from the reaction at the indicated time points and target cleavage was terminated by adding EDTA to a final concentration of 50 mM. Cleavage products at a 4 nM final concentration were separated by capillary electrophoresis (CE) on an Applied Biosystems 3730xl DNA Analyzer (Applied Biosystems, Waltham, MA, USA). The quantitative analysis of fluorescent peaks was performed using PeakScanner Software v1.0 (Thermo Fisher Scientific, Inc., Waltham, Massachusetts, USA) and fragment analysis software for in-house use at New England Biolabs as previously described (29, 43).

### Argonaute cleavage assays on double-stranded DNA

A segment of phage φX174 DNA (3536-3858 nt) was amplified by PCR to generate a 5’-FAM/ROX labeled 322 bp DNA substrate. The PCR product was purified using Monarch PCR and DNA Cleanup kit (New England Biolabs, Inc., Ipswich, MA, USA) and DNA concentration was quantified using NanoDrop spectrophotometer (Thermo Fisher Scientific, Inc., Waltham, Massachusetts, USA). For single-strand cleavage experiments the CbAgo/guide complex was formed by incubating 0.5 μM CbAgo and 1 μM guide for 15 minutes at 37°C in a 10 μl of 1X CutSmart buffer (50 mM Potassium Acetate, pH 7.9, 20 mM Tris-Acetate, 10 mM Magnesium Acetate, 100 μg/ml BSA). The CbAgo/guide complex at a 0.25/0.5 μM final concentration was combined with 50 nM 322 bp 5’-FAM/ROX labeled DNA substrate and 0.5 μM RecB^exo-^C in 20 μl of 1X CutSmart buffer and cleavage reaction was initiated at 37°C by adding 5 mM ATP. For double-strand cleavage experiments, two separate 5 μl reactions, each containing CbAgo (0.5 μM) and guide (1 μM), were carried out to form CbAgo/guide complexes that target opposing DNA strands. CbAgo/guide complexes, each at a 0.125/0.25 μM final concentration, were combined with 50 nM 322 bp 5’FAM/ROX labeled DNA substrate and 0.25 μM RecB^exo-^C in 20 μl of 1X CutSmart buffer and reaction was initiated at 37°C by adding 5 mM ATP. The cleavage products were analyzed by CE as described above. For time course experiments the cleavage reaction volume was increased to 50 μl and 5 μl samples were withdrawn from the reaction at the indicated time points. Cleavage of φX174, pAd2_BsaI and pAd2_AvrII was carried out using DNA linearized with the appropriate restriction enzyme. Reactions contained 0.2 μg of linear DNA, 5mM ATP and two CbAgo/guide complexes, each at 0.125/0.25 μM concentration, in 20 μl of 1X CutSmart buffer. The CbAgo cleavage was carried out at 37ºC for 1h either in the absence or in the presence of 0.5 μM RecB^exo-^C. The reactions were stopped by treatment with 8 units of Proteinase K for 30 minutes at 25ºC. CbAgo cleavage products were column purified and separated by gel electrophoresis on 1.2% agarose gel. The plasmids pAd2_BsaI and pAd2_ AvrII were created by Dr. Richard Morgan (New England Biolabs, Inc., Ipswich, MA, USA) by cloning either a 22404 bp BsaBI fragment or 19428 bp AvrII fragment of Adenovirus2 genomic DNA into pUC19 vector.

### Ligation of CbAgo cleaved DNA fragments

A 332bp 5’-FAM/ROX labeled DNA1 was generated by PCR using a segment of φX174 DNA (nucleotides 3681-4012) as a template. DNA1 was cleaved with CbAgo loaded with guides T2+B1 to generate a 15 bp 5’-ROX labeled and a 309 bp 5’-FAM labeled cleavage products. A 300 bp 5’-FAM/ROX labeled DNA2 was generated by PCR using a segment of pUC19 DNA (nucleotides 806-1105) as a template. DNA2 was cleaved with CbAgo loaded with guides T5+B5 to generate a 14 bp 5’-ROX labeled and a 278 bp 5’-FAM labeled cleavage products. All CbAgo cleavage products carried 8-nt long 3’-ss overhangs. The enzymes were inactivated by treatment with Proteinase K for 30 min at 37ºC. The 15 bp and 14 bp fragments were discarded by column purification using Monarch PCR and DNA Cleanup Kit. A 29 bp ds “Bridge” oligonucleotide was created by combining two complementary 5’-phophorylated ssDNA oligonucleotides (1 nmol each) in 100 μl of 10 mM Tris-HCl buffer, pH 7.5 and heating for 5 min at 95ºC followed by slow cooling down to room temperature. On both ends, the Bridge oligo carried 8-nt long 3’-ss overhangs complementary to the 3’-ss overhangs on the 309 bp and 278 bp CbAgo cleavage products. DNA fragment ligation was carried out with T4 DNA ligase (400 units) in 10 μl of T4 DNA ligase buffer. First, 0.3 pmols of 309 bp DNA1 fragment and 0.3 pmols of 29 bp bridge oligo were ligated for 15 minutes at 25ºC. The reaction was then supplemented with 0.3 pmols of 278 bp DNA2 fragment and ligation continued for another 15 minutes at 25ºC. Ligation products were analyzed by CE.

## ACKNOWLEDGEMENTS

The authors thank Drs. W. Jack, A. Gardner, N. Tanner and E. Hunt for helpful discussions and critical comments on the manuscript; Dr. R. Morgan for providing DNA substrate plasmids; Dr. P. Hsieh and J.Greci for assistance with RecBC constructs and purification; L. Mazzola, D. Fuchs, K. Augulewicz and H. Bell for performing core laboratory capillary electrophoresis analysis. This study is supported by New England Biolabs, Inc.

## Author contributions

J.B. designed research; R.V., J.V., J.M. and J.B. performed research; V.P. performed bioinformatics analysis; J.B., R.V and V.P wrote the paper.

## SUPPLEMENTARY INFORMATION

### Supplementary Methods

#### pAgo protein expression and purification

T7 Express lysY/I^q^ *E*.*coli* (New England Biolabs) carrying expression plasmid was grown at 37°C in 1L of LB medium containing 0.02 mg/ml kanamycin until OD_600_ 0.6. Protein expression was induced with 0.2 mM IPTG and cell culture continued to grow overnight at 16°C. The cells were harvested by centrifugation, resuspended in 45 ml buffer A (20 mM Tris-HCl, pH 7.5, 300 mM NaCl, 10 mM imidazole) supplemented with protease inhibitor cocktail (Complete mini, EDTA-free; Roche diagnostics) and disrupted by sonication. The cell-free lysate was loaded on a 5 ml HisTrap Nickel HP column (GE-Healthcare) and proteins were eluted with 20-250 mM imidazole gradient. The fractions containing Ago protein were pooled, diluted 2-fold with buffer B (20 mM Tris-HCl, pH 7.5, 0.1 mM EDTA), loaded on 5 ml HiTrap Heparin HP column (GE-Healthcare) and proteins were eluted with 0.15-1 M NaCl gradient. The fractions containing Ago protein were pooled, diluted with buffer B to 150 mM NaCl concentration, loaded on 5 ml HiTrap Capto S Sepharose HP column (GE-Healthcare) and eluted with 0.15-0.5 M NaCl gradient. The fractions containing Ago were pooled and loaded on 5 ml Bioscale CHTP column (Bio-Rad). The proteins were eluted with 0.02-0.3 M KPO4 gradient. The purified protein was concentrated by dialysis against 20 mM Tris-HCl (pH 7.5), 300 mM NaCl, 0.1 mM EDTA, 0.1 mM TCEP and 50% (vol/vol) glycerol and stored at −80°C. The purified argonaute proteins were >95% homogenous as determined by Coomassie-stained SDS-PAGE.

#### RecB protein purification

T7 Express *E*.*coli* (New England Biolabs) carrying RecB^exo-^ expression plasmid was grown at 37°C in until OD_600_ =0.6. Protein expression was induced with 0.2 mM IPTG and cell culture continued to grow overnight at 16°C. The cells were harvested by centrifugation, resuspended in 150 ml buffer A (20 mM potassium phosphate, pH 7.0, 300 mM NaCl, 20 mM imidazole, 5% glycerol) and disrupted by sonication. The clarified lysate was loaded on a HisTrap Nickel HP column (5 ml) and proteins were eluted using 20-250 mM imidazole gradient in buffer A. The fractions containing RecB^exo-^ protein were diluted 2-fold with buffer B (20 mM potassium phosphate, pH 7.0, 0.1 mM EDTA, 5% glycerol) to bring NaCl concentration to 100 mM and loaded on HiTrap Heparin HP column (5 ml). The proteins were eluted using 0.1-0.8 M NaCl gradient in buffer B. The fractions containing RecB^exo-^ protein were diluted 2-fold with buffer B and loaded on Bioscale CHTP column (5 ml). The proteins were eluted with 0.02-0.4 M KPO4 gradient in buffer B containing 250mM NaCl. The purified protein was concentrated by dialysis against 10 mM Tris-HCl, pH 7.4, 300 mM NaCl, 0.1 mM EDTA, 0.1 mMDTT and 50% glycerol. The protein was >95% homogenous as determined by Coomassie-stained SDS-PAGE.

#### RecC protein purification

T7 Express *E*.*coli* carrying RecC expression plasmid was grown and expressed as described above for RecB^exo-^ protein. The cells were resuspended in 80 ml of 20 mM potassium phosphate buffer, pH 7.0 containing 450 mM NaCl, 20 mM imidazole and disrupted by sonication. The clarified lysate was loaded on a HisTrap Nickel HP column (5 ml) and proteins were eluted with 20-250 mM imidazole gradient in the buffer that contained 300 mM NaCl. The pooled RecC protein was diluted 3-fold with 20 mM potassium phosphate buffer, pH 7.0, 0.1 mM EDTA, 5% glycerol to bring NaCl concentration to 100 mM and was loaded on HiTrap Heparin HP column (5 ml). RecC protein did not bind to the Heparin column at 100 mM NaCl. The flow through fraction was further diluted to 75 mM NaCl concentration and loaded on HiTrap Q HP column (5ml). RecC protein was eluted with 0.75-1.0 M NaCl gradient and concentrated by dialysis against 10 mM Tris-HCl, pH 7.4, 300 mM NaCl, 0.1 mM EDTA, 0.1 mM DTT and 50% glycerol. The protein was >90% homogenous as determined by Coomassie-stained SDS-PAGE.

#### Helicase assay for DNA unwinding activity

We have modified the previously reported gel-electrophoresis assay for monitoring helicase activity (1) to run with a high-throughput CE technique. We employed restriction endonuclease DpnI which cleaves a fully methylated G^m^ATC/G^m^ATC site but does not cleave a hemimethylated site (G^m^ATC/GATC). Helicase assay uses a 5’-FAM labeled DNA substrate comprised of 30 bp duplex portion and 30-nt long fork structure. 5’-FAM labeled strand has a G^m^ATC sequence within the duplex portion, whereas the opposite strand has an unmethylated GATC site (Supplementary Figure S2A). The ds fork substrate is mixed at a 1:50 molar ratio with 30-nt long ss trap oligo which has a methylated G^m^ATC site and is complementary to the 5’-FAM labeled strand of DNA substrate. The substrate/trap mixture is then incubated with DNA helicase and DpnI. As soon as helicase unwinds ds substrate, the trap oligo anneals to the methylated strand creating a hybrid duplex with a site G^m^ATC/G^m^ATC which is immediately cleaved by DpnI. The progress of DNA strand unwinding can be monitored by the disappearance of a 5’-FAM labeled substrate and by the accumulation of a 5’-FAM labeled DpnI cleavage product, both visualized as different size peaks after resolving them by CE. DNA oligonucleotides carrying ^m6^A residue within a GATC site were synthesized at Organic Synthesis facility of New England Biolabs. Double-stranded fork substrate was created by annealing two complementary ss oligonucleotides (Supplementary Table S1). DNA unwinding reactions were carried out for 15 minutes at 37ºC in 10 μl of 1X CutSmart buffer containing 0.1 μM fork substrate, 5 μM trap oligo, 5 mM ATP, 2 units DpnI and 1 μl of serially diluted DNA helicase. Reactions were terminated by adding 50 mM EDTA. Cleavage products were analyzed by CE as described above.

## SUPPLEMENTARY TABLES

**Supplementary Table S1.**
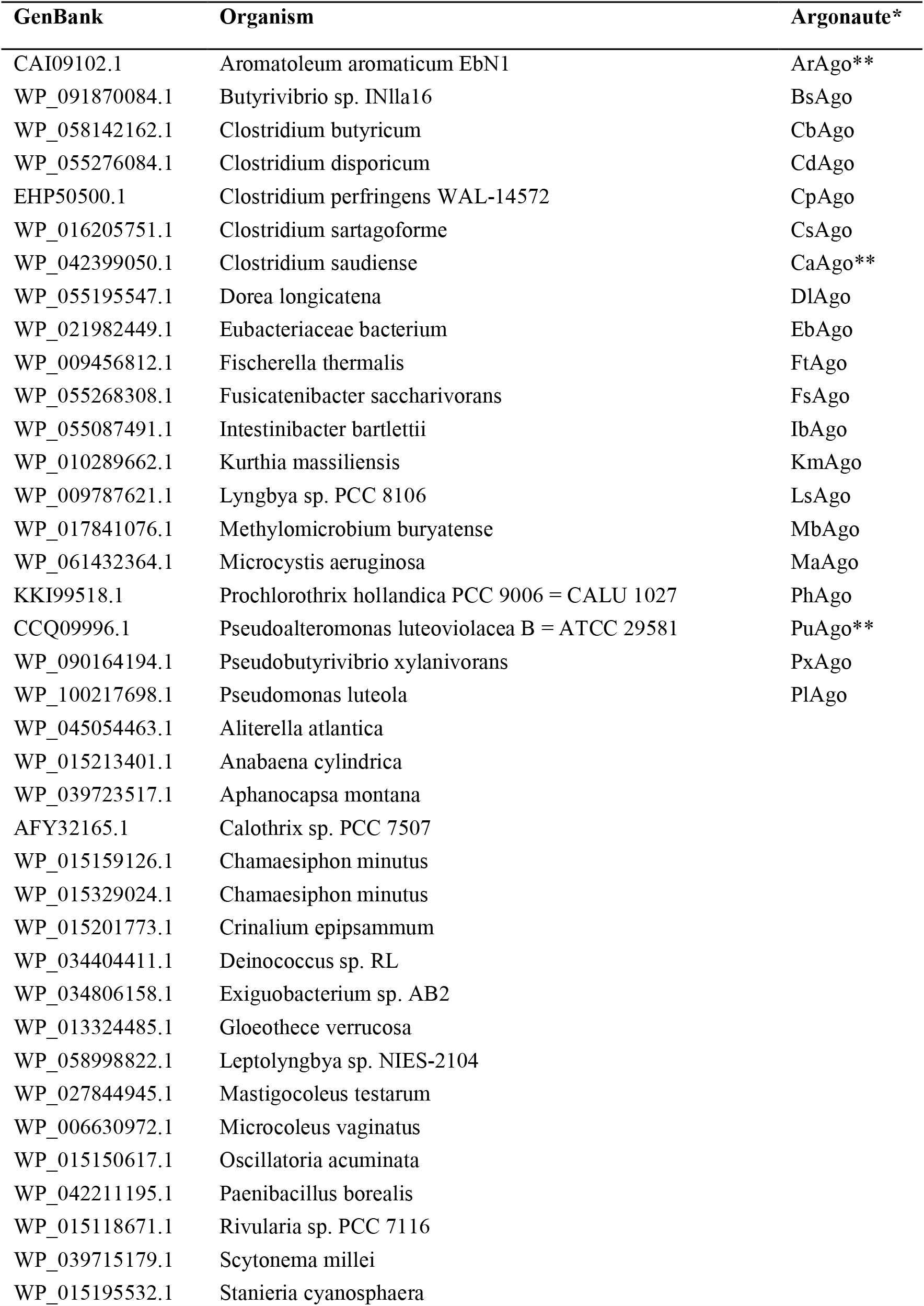
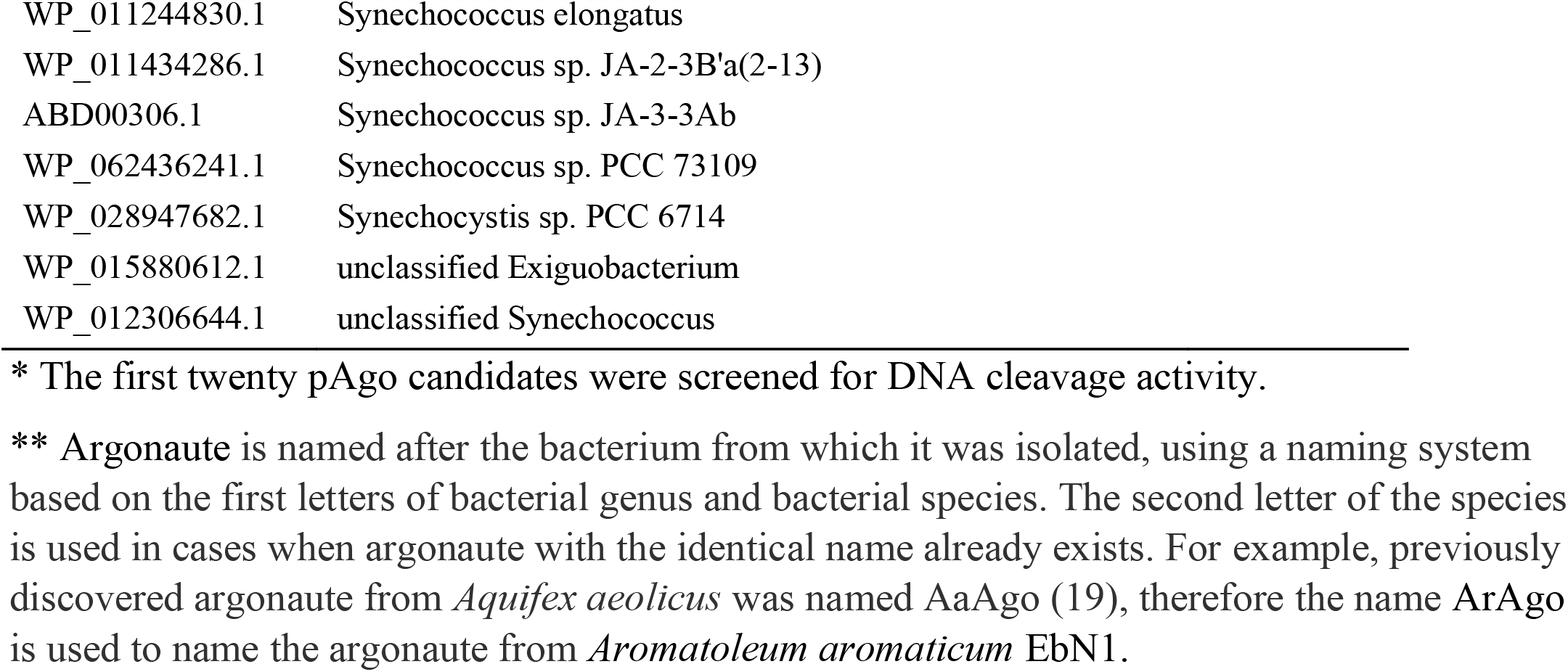
The list of argonaute proteins.

**Supplementary Table S2.** Cleavage efficiency of pAgo candidates.

| Species | Argonaute | DNA-Guided Cleavage of |  | RNA-Guided Cleavage of |  |
| --- | --- | --- | --- | --- | --- |
|  |  | DNA, % | RNA, % | DNA, % | RNA, % |
| *Clostridium butyricum* | CbAgo | 100 | 0 | 0 | 0 |
| *Clostridium perfringens* | Cp Ago | 96.6±0.8 | 61.7±3.8 | 18.5±1.0 | 0 |
| *Clostridium saudiense* | Ca Ago | 83.0±2.2 | 0 | 0 | 0 |
| *Intestinibacter bartlettii* | IbAgo | 68.5±2.4 | 0 | 0 | 0 |
| *Clostridium sartagoforme* | Cs Ago | 65.2±4.3 | 0 | 0 | 0 |
| *Clostridium disporicum* | Cd Ago | 63.5±3.9 | 0 | 0 | 0 |

Cleavage of either target T-2 (DNA) or target T-3 (RNA) was carried out with the purified pAgo proteins using either guide G-2 (DNA) or G-3 (RNA). Cleavage reactions were performed at Ago:guide:target concentration ratio of 250:250:50 nM, and reactions were incubated for 1 hour at 37°C. Data represents means and standard deviations from three independent experiments.

**Supplementary Table S3.**
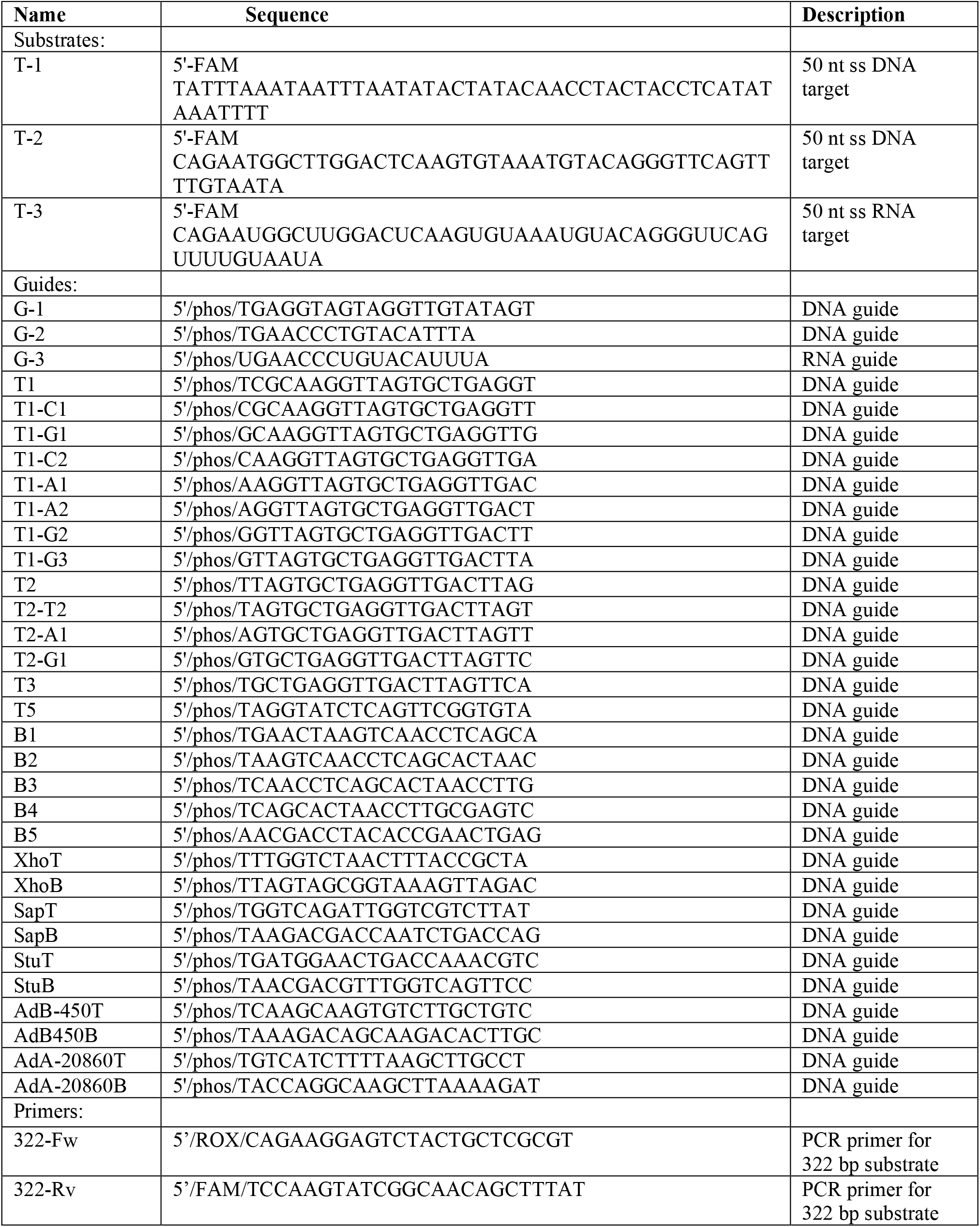
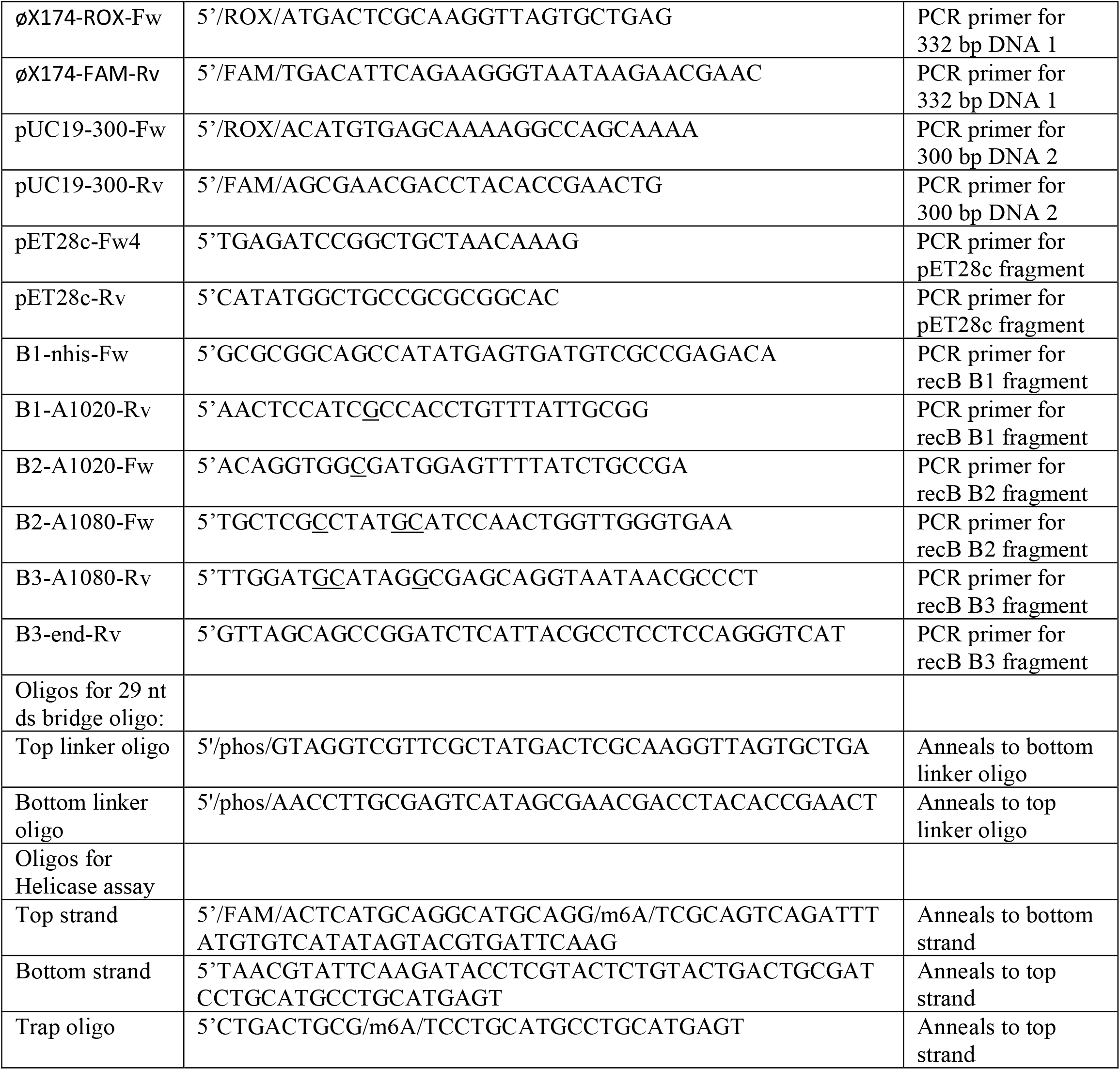
Oligonucleotide sequence used in this work. Nucleotides in PCR primers that were changed to introduce mutations in the recB gene are underlined.

## SUPPLEMENTARY FIGURES

**Supplementary Figure S1.**
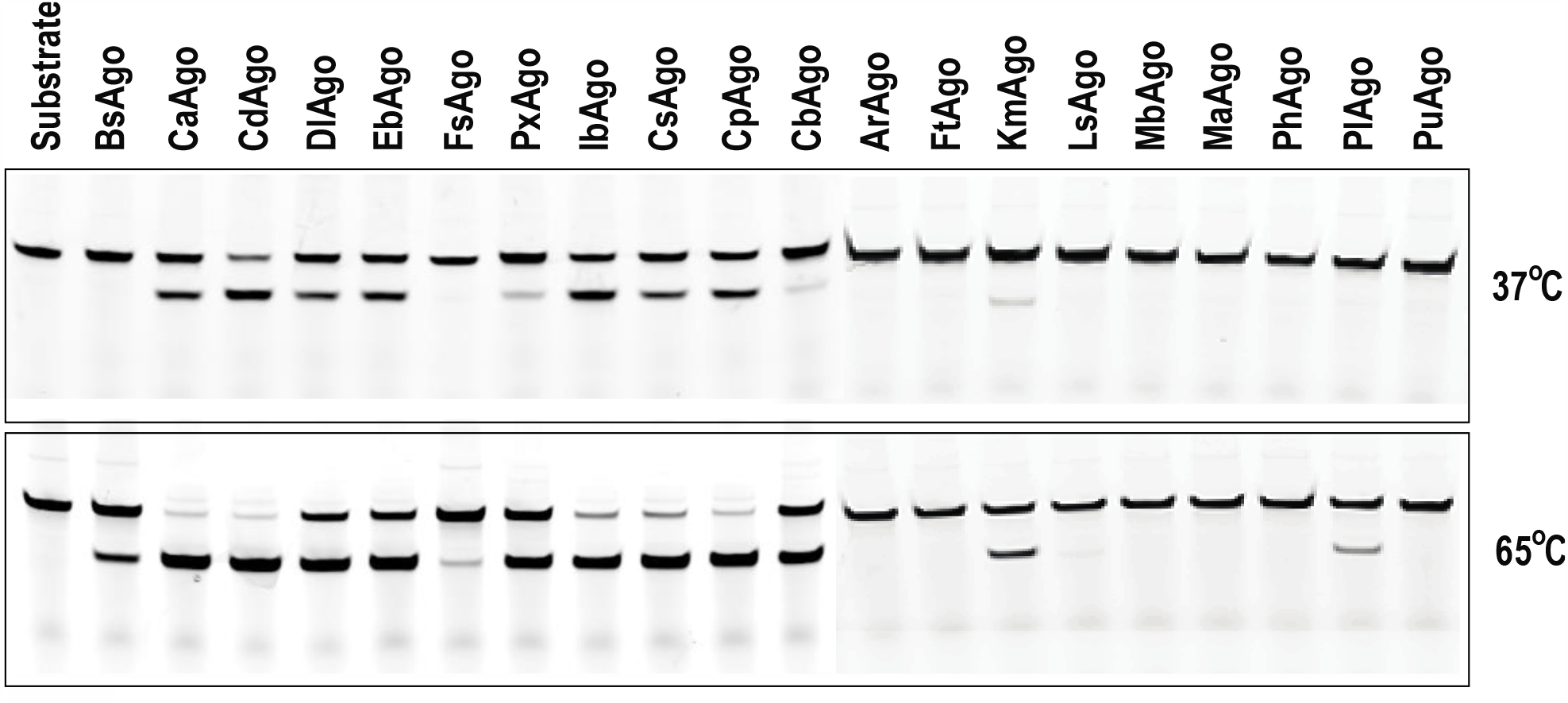
Putative Argonaute candidates were tested for DNA-guided cleavage activity at 37°C and 65°C. Argonaute proteins were synthesized *in vitro* using PURExpress protein synthesis kit. ssDNA target was cleaved with the indicated pAgo for 1 hour at either 37ºC or 65ºC. Cleavage products were separated by electrophoresis on denaturing polyacrylamide gel and 5’-FAM labeled products were visualized with Typhoon 9400 Scanner. Candidate pAgo names are shown above the respective lane. Bacterial hosts and NCBI Gene bank Accession numbers are listed in Supplementary Table S1.

**Supplementary Figure S2.**
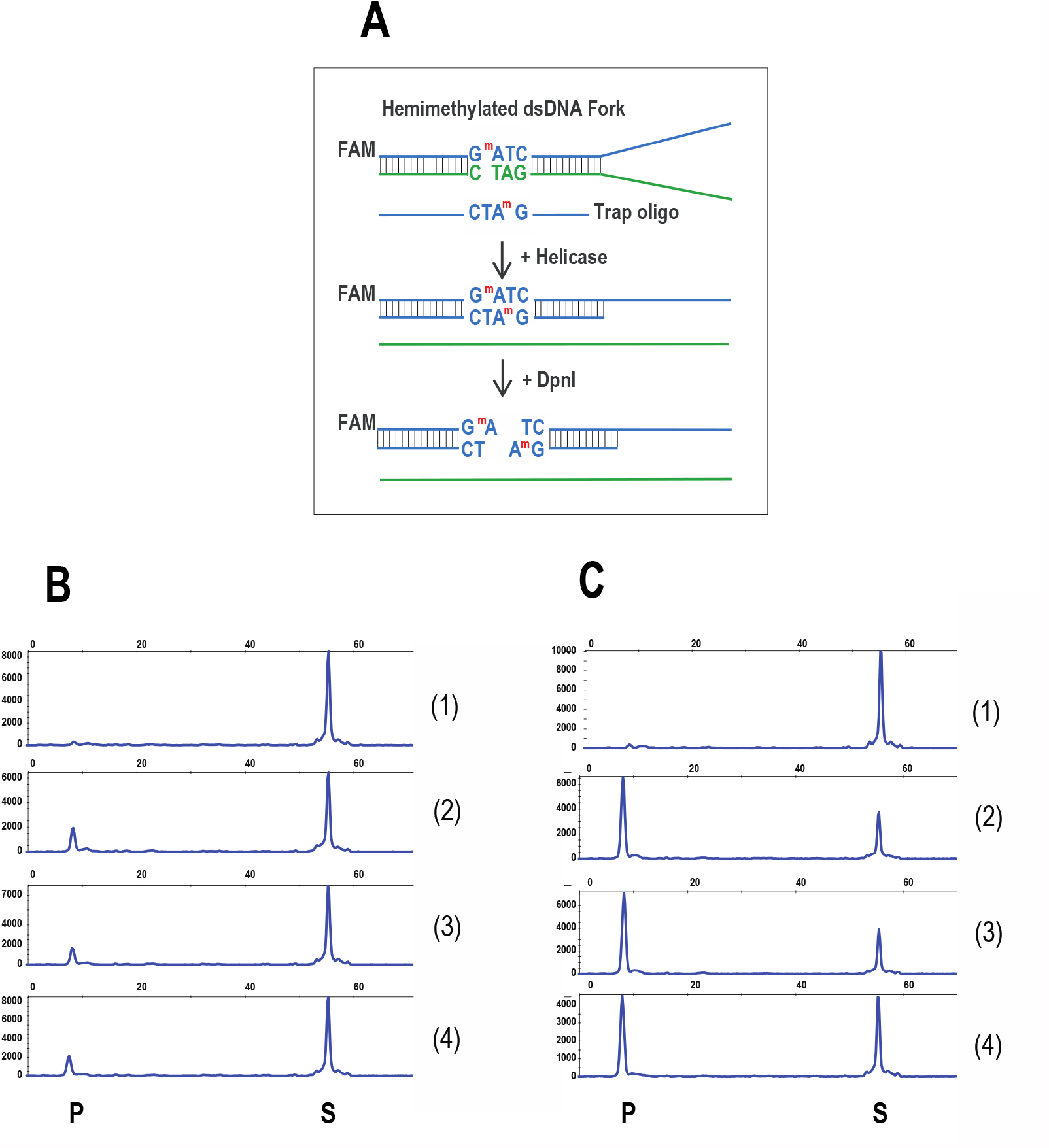
DNA unwinding activity of RecB^exo-^ and RecB^exo-^C DNA helicases. **(A)** Schematic overview of helicase activity assay using DpnI restriction endonuclease. RecB^exo-^ and RecB^exo-^C DNA helicases were serially diluted, and 1 μl of serially diluted protein was added to 10 μl of 1X CutSmart buffer containing 0.1 μM fork substrate, 5 μM trap oligo, 5 mM ATP and 2 units DpnI. DNA unwinding reactions were carried out for 15 minutes at 37ºC and were terminated by adding 50 mM EDTA to the final concentration. Cleavage products were analyzed by CE. **(B)** Reactions were carried out in the absence (lane 1) or in the presence of 2.5, 1.25 and 0.63 μM RecB^exo-^ (lanes 2-4, respectively). **(C)** Reactions were carried out in the absence (lane 1) or in the presence of 1.0, 0.5 and 0.25 μM RecB^exo-^C (lanes 2-4, respectively). S - DNA substrate; P-DpnI cleavage product.

**Supplementary Figure S3.**
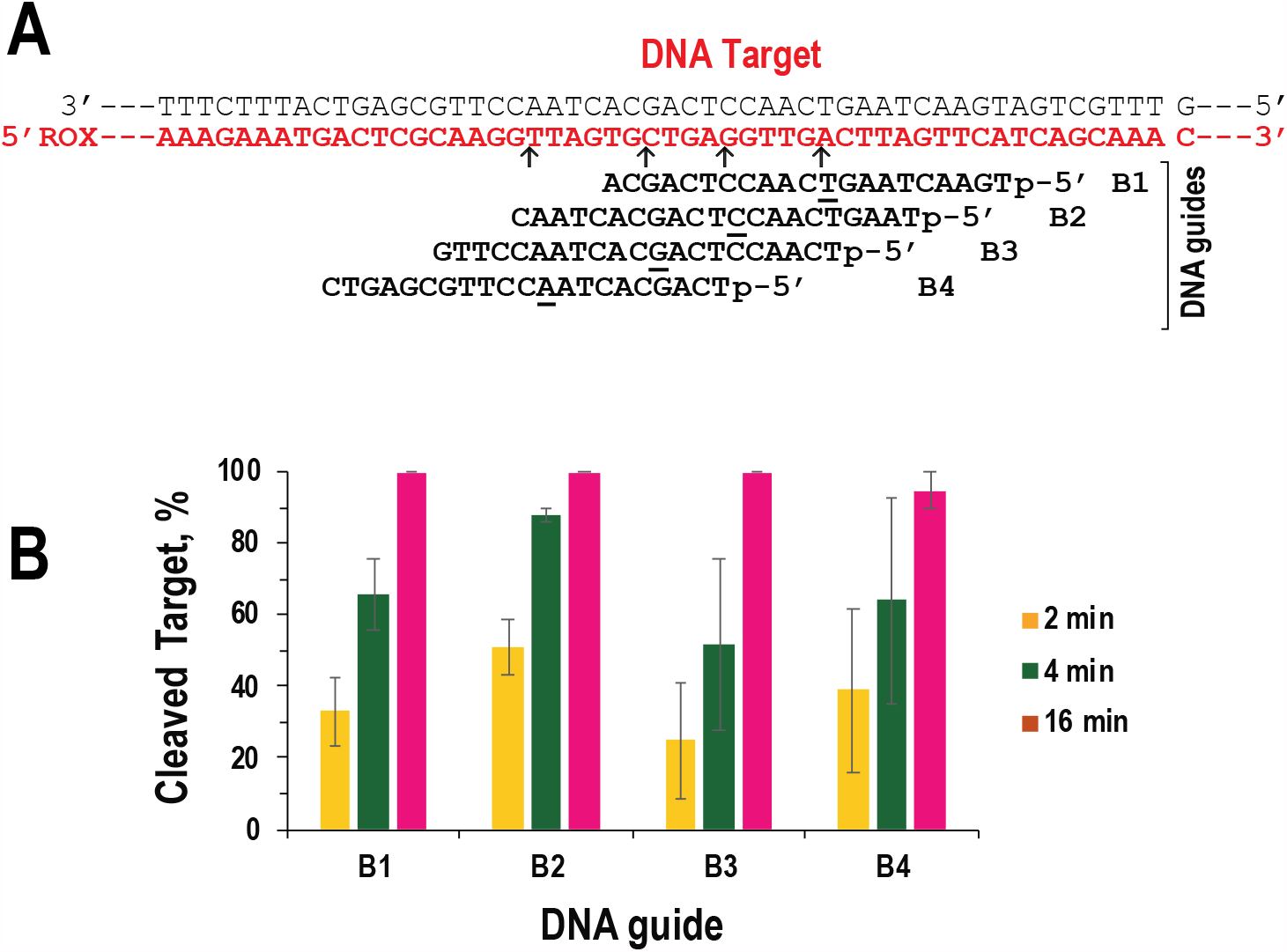
CbAgo cleavage of 5’-ROX labeled DNA strand after dsDNA is unwound by RecB^exo-^C helicase. (**A**) Arrangement of DNA guides on 322 bp dsDNA substrate. 5’-phosphorylated guides are complementary to the 5’-ROX labeled strand (shown in red). Within each guide sequence, the 10^th^ nucleotide starting from a 5’-phosphate is underlined to mark position that aligns with the CbAgo cut site (shown by an arrow). (**B**) Strand cleavage efficiency by CbAgo loaded with the indicated guides. Reactions were carried out at CbAgo:guide:target molar concentration ratio of 250:500:50 nM. The percentage of cleaved DNA target was quantified after 2, 4 and 16 minutes at 37°C. Error bars indicate standard deviation from three independent experiments.

**Supplementary Figure S4.**
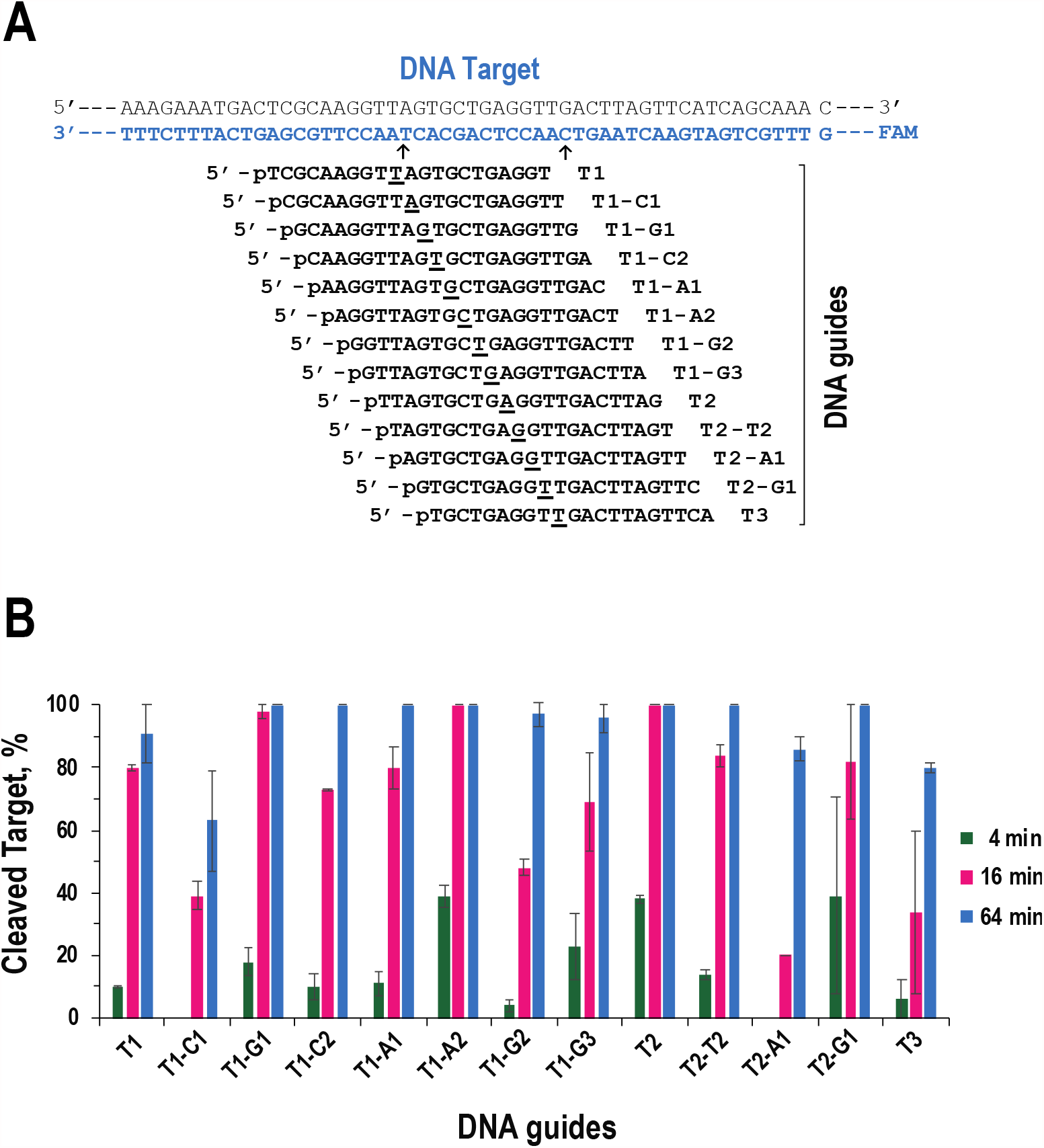
CbAgo cleavage of 5’-FAM labeled DNA strand after dsDNA is unwound by RecB^exo-^C helicase. (**A**) Arrangement of DNA guides on 322 bp dsDNA substrate. ssDNA guides are complementary to the 5’-FAM labeled strand (shown in blue). Arrow on the left shows the cleavage position of CbAgo loaded with a guide T1. Arrow on the right shows the cleavage position of CbAgo loaded with a guide T3. (**B**) Strand cleavage efficiency by CbAgo loaded with the indicated guides. Reactions were carried out at CbAgo:guide:target molar concentration ratio of 250:500:50 nM. The percentage of cleaved DNA target was quantified after 4, 16 and 64 minutes at 37°C. Error bars indicate standard deviation of two independent experiments.

**Supplementary Figure S5.**
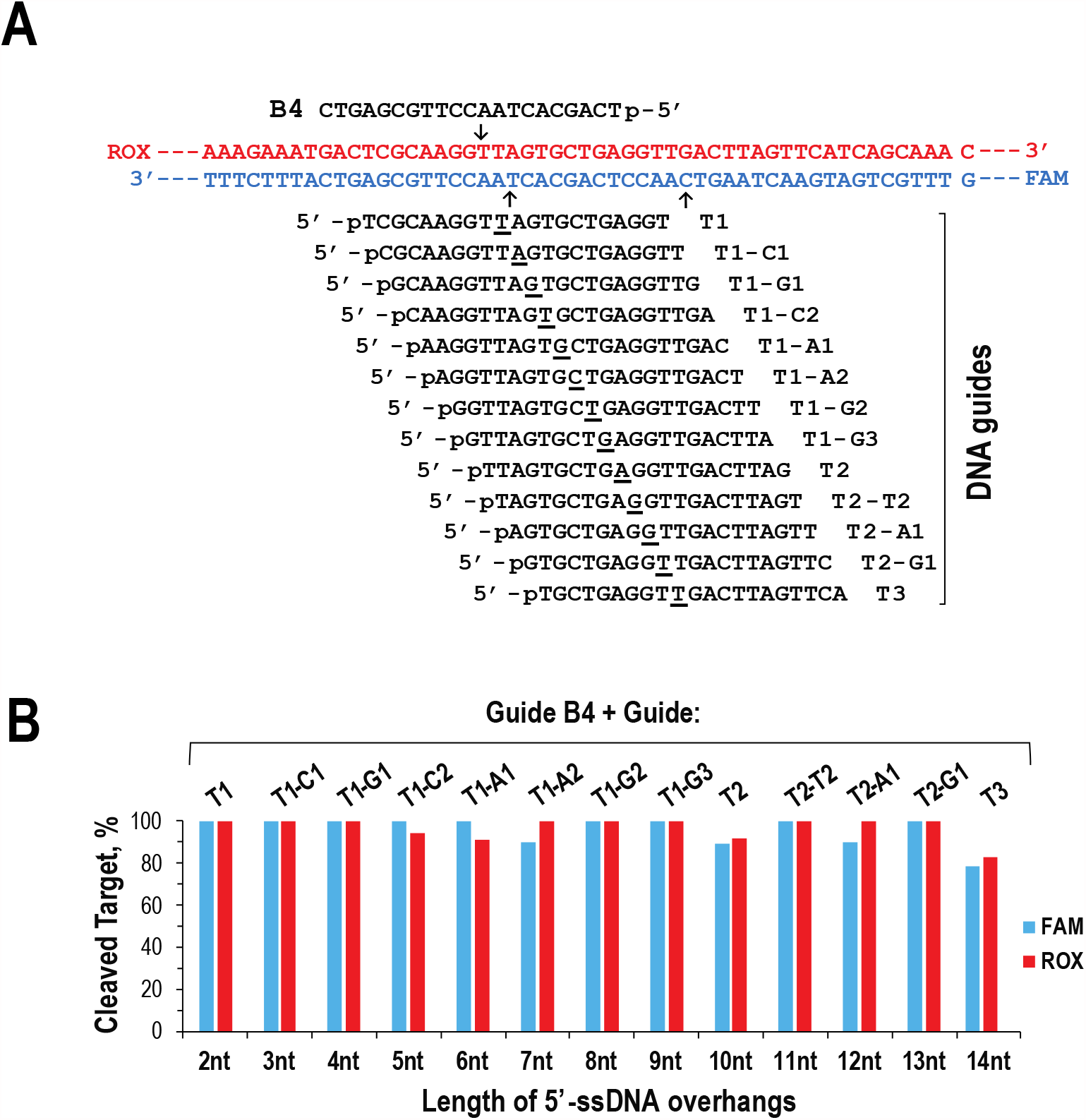
dsDNA cleavage at a 1:1 CbAgo:guide molar ratio. **(A)** Schematic overview of guide positioning on dsDNA target. 5’-ROX labeled DNA strand is cleaved by CbAgo/B4 complex. 5’-FAM labeled DNA strand is targeted with thirteen guides which are shifted by one nucleotide with respect to each other along DNA strand. Arrow on the left indicates the cleavage position of CbAgo/T1 complex. Arrow on the right indicates the cleavage position of CbAgo/T3 complex. **(B)** Efficiency of double-strand cleavage by CbAgo loaded with the indicated guide pairs which generate cleavage fragments tailed with 5’-ss overhangs varying from 2 to 14 nt in length. CbAgo/guide complexes were combined with DNA target at CbAgo:guide:target molar ratio of 125:125:50 nM. The concurrent double-strand cleavage was carried out for 32 minutes at 37ºC in the presence of 250 nM RecB^exo-^C. The percentage of cleaved DNA was quantified for each DNA strand.

**Supplementary Figure S6.**
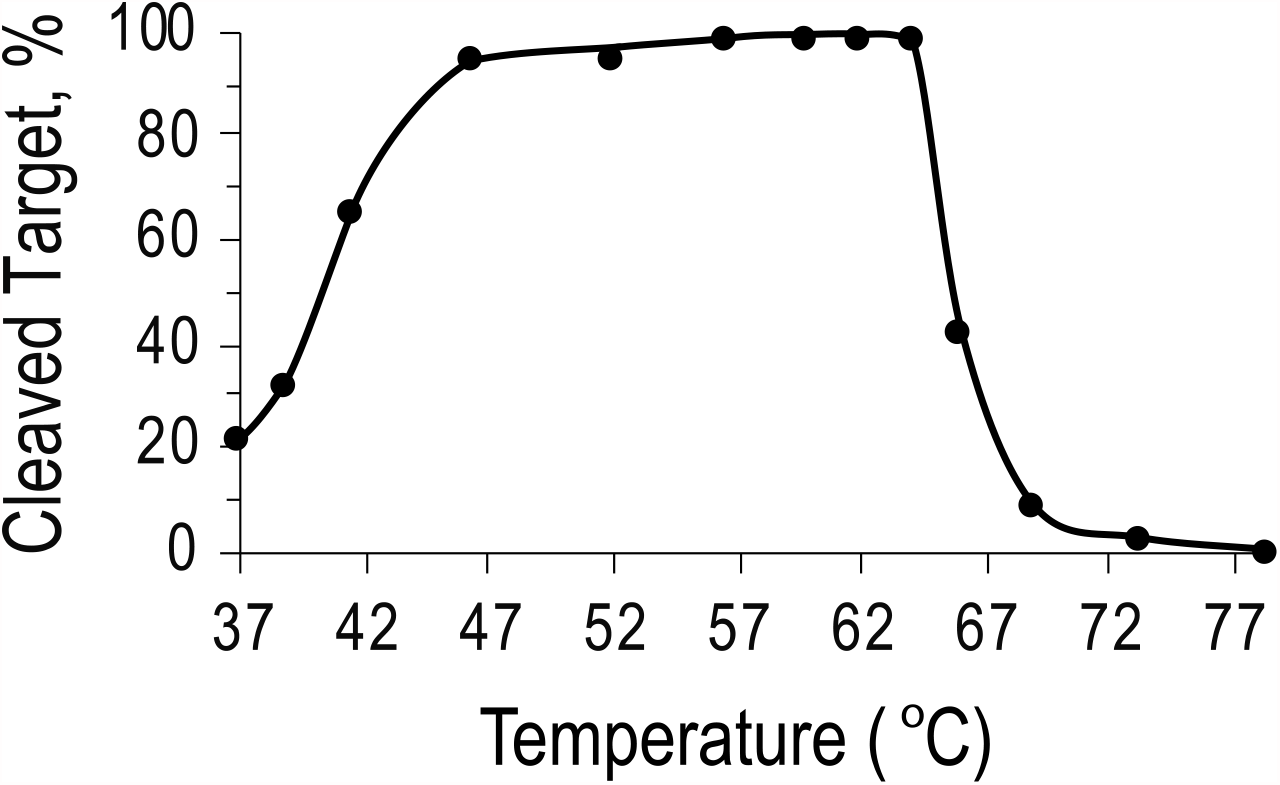
CbAgo activity at different reaction temperatures. CbAgo was loaded with guide G-2 at 37°C. 5’-FAM labeled single-stranded substrate T-2 was cleaved for 10 minutes at indicated temperature at CbAgo:guide:target concentration ratio of 15:30:50 nM. Cleavage products were analyzed by CE.

**Supplementary Figure S7.**
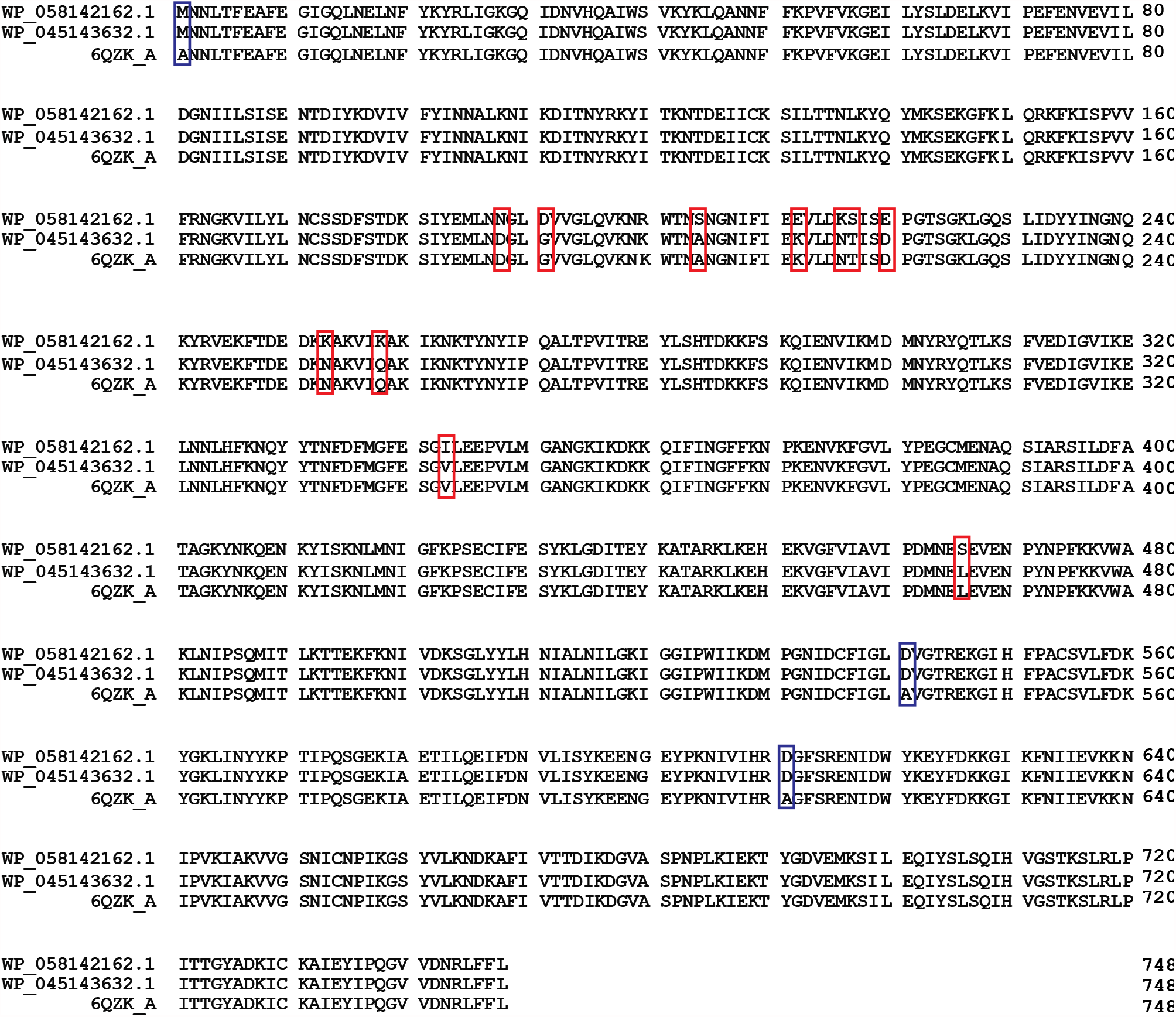
Amino acid sequence alignment of CbAgo variants studied by different research groups. WP_058142162.1 shows a.a. sequence of CbAgo studied by us and by group referred in (24), WP_045143632.1 shows a.a. sequence of CbAgo referred in (42) and 6QZK_A shows a.a. sequence of CbAgo referred in (30). Red boxes indicated different amino acids; blue boxes indicate amino acid substitutions which were introduced by authors to make the CbAgoDM variant for structural studies (30).

## REFERENCES

1. L. Peters and G. Meister, Argonaute proteins: mediators of RNA silencing. Mol. Cell., 26, 611–623 (2007).

2. S.M. Hammond, E. Bernstein, D. Beach, C.J. Hannon, An RNA directed nuclease mediated post-transcriptional gene silencing in Drosophila cells. Nature, 404, 293–296 (2000).

3. G. Hutvagner, M.J. Simard, Argonaute proteins: key players in RNA silencing. Nat. Rev. Mol. Cell Biol. 9, 22–32 (2008).

4. M. Ghildiyal, P.D. Zamore, Small silencing RNAs: an expanding universe. Nat. Rev. Genet.,10, 94–108 (2009).

5. J. Martinez, A. Patkaniowska, H. Urlaub, R. Luhrmann, T. Tuschl, Single-Stranded Antisense siRNAs Guide Target RNA Cleavage in RNAi introduced dsRNA. Cell, 110, 563–574 (2002).

6. S.M. Hammond, E. Bernstein, D. Beach, G.J. Hannon, An RNA-directed nuclease mediates post-transcriptional gene silencing in Drosophila cells. Nature, 404, 293–296 (2000).

7. S.M. Elbashir, J. Harborth, W. Lendeckel, A. Yalcin, K. Weber, et al. Duplexes of 21-nucleotide RNAs mediate RNA interference in mammalian cell culture. Nature, 411, 494–498 (2001).

8. A. J. Pratt, I. J. MacRae, The RNA-induced silencing complex: a versatile gene-silencing machine. J. Biol. Chem., 284, 17897–17901 (2009).

9. D. Moazed, Small RNAs in transcriptional gene silencing and genome defence. Nature, 457, 413–420 (2009).

10. K.S. Makarova, Y.I. Wolf, J. van der Oost, E.V. Koonin, Prokaryotic homologs of Argonaute proteins are predicted to function as key components of a novel system of defense against mobile genetic elements. Biol. Direct., 4, 29 (2009).

11. I. Olovnikov, K. Chan, R. Sachidanandam, D. Newman, A. Aravin, Bacterial Argonaute Samples the transcriptome to identify Foreign DNA. Cell, 51, 594–605 (2013).

12. D.C. Swarts, K. Makarova, Y. Wang, K. Nakanishi, R.F. Ketting, et al. The evolutionary journey of Argonaute proteins. Nat. Struct. Mol. Biol., 21, 743–753 (2014).

13. K. Hur, I. Olovnikov, A.A. Aravin, Prokaryotic argonautes defend genomes against invasive DNA. Trends Biochem. Sci., 39, 257–259 (2014).

14. A. Kuzmenko, A. Oguienko, D. Esyunina, D. Yudin, M. Petrova, et al. DNA targeting and interference by a bacterial Argonaute nuclease. Nature, 587, 632–637 (2020).

15. S. Ryazansky, A. Kulbachinskiy, A.A. Aravin, The Expanded Universe of Prokaryotic Argonaute Proteins, mBio., 9, e01935–18 (2018).

16. Y. Wang, S. Juranek, H. Li, G. Sheng, G.S. Wardle, et al. Nucleation, propagation and cleavage of target RNAs in argonaute silencing complexes. Nature, 461, 754–761 (2009).

17. G. Sheng, H. Zhao, J. Wang,Y Rao, W. Tian, et al. Structure-based cleavage mechanism of Thermus thermophilus Argonaute DNA guide strand-mediated DNA target cleavage. Proc. Natl. Acad. Sci. USA, 111, 652–657 (2014).

18. K.W. Doxzen, J.A. Doudna, DNA recognition by an RNA-guided bacterial Argonaute. PLoS One, 12, e0177097 (2017).

19. Y.R. Yuan, Y. Pei, J.B. Ma, V. Kuryavyi, M. Zhadina et al. Crystal structure of A. aeolicus argonaute, a site-specific DNA-guided endoribonuclease, provides insights into RISC-mediated mRNA cleavage. Mol. Cell., 19, 405–419 (2005).

20. D.C. Swarts, J.W. Hegge, I. Hinojo, M. Shiimori, M.A. Ellis et al. Argonaute of the archaeon Pyrococcus furiosus is a DNA-guided nuclease that targets cognate DNA. Nucl. Acids Res., 43, 5120–5129 (2015).

21. A. Zander, P. Hozmeister, D. Klose,P. Tinnefeld, D. Grohmann, Single-molecule FRET supports the two-state model of Argonaute action. RNA Biol., 11, 45–56 (2014).

22. D.C. Swarts, M. Szczepaniak, G. Sheng, S.D. Chandradoss, Y. Zhu et al. Autonomous Generation and Loading of DNA Guides by Bacterial Argonaute. Mol. Cell., 65, 985–998 (2017).

23. A. Zander, S. Willkomm, S. Ofer, M. van Wolferen, L. Egert et al. Guide-independent DNA cleavage by archaeal Argonaute from Methanocaldococcus jannaschii. Nat. Microbiol., 2, 1–10 (2017).

24. A. Kuzmenko, D. Yudin, S. Ryazansky, A. Kulbachinskiy, A.A. Aravin, Programmable DNA cleavage by Ago nucleases from mesophilic bacteria Clostridium butyricum and Limnothrix rosea. Nucl. Acids Res., 47, 5822–5836 (2019).

25. D.C. Swarts, M.M. Jore, E.R. Westra, Y. Zhu, J.H. Janssen et al. DNA-guided DNA interference by a prokaryotic Argonaute. Nature, 507, 258–261 (2014.)

26. J.W. Hegge, D.C. Swarts, J. van der Oost, Prokaryotic Argonaute proteins: novel genome-editing tools? Nat. Rev. Microbiol.,16, 5–11 (2018).

27. H. Wang, M. La Russa, L.S. Qi, CRISPR/Cas9 in Genome Editing and Beyond. Annu. Rev. Biochem., 85, 227–264 (2016).

28. B. Enghiad, H. Zhao, Programmable DNA-guided Artificial Restriction Enzymes. ACS Synthetic Biology. ACS Synth. Biol., 6, 752–757 (2017).

29. E.A., Hunt, T.C. Evans Jr., N.A. Tanner, Single-stranded binding proteins and helicase enhance the activity of prokaryotic argonautes in vitro. PLoS One, 13, e0203073 (2018).

30. J.W. Hegge, D.C. Swarts, S.D. Chandradoss, T.J. Cui, J. Kneppers et al. DNA-guided DNA cleavage at moderate temperatures by Clostridium butyricum Argonaute. Nucl. Acids Res., 47, 5809–5821 (2019).

31. Y. Cao, W. Sun, J. Wang, G. Sheng, G. Xiang et al. Argonaute proteins from human gastrointestinal bacteria catalyze DNA-guided cleavage of single-and double-stranded DNA at 37°C. Cell Discov., 5, 1–4 (2019).

32. A. Olina, A. Kuzmenko, M. Ninova, A.A. Aravin, A. Kulbachinskiy et al. Genome-wide DNA sampling by Ago nuclease from the cyanobacterium Synechococcus elongatus. RNA Biol.,17, 677–688 (2020).

33. S. Willkomm, A. Zander, A. Gust, D.A. Grohmann, Prokaryotic twist on argonaute function. Life, 5, 538–553 (2015).

34. K.D. Raney, A.K. Byrd and S. Aarattuthodiyil, Structure and mechanism of SF1 DNA helicases. Adv. Exp. Med. Biol., 767, 17–46 (2013).

35. A.F. Taylor, G.R. Smith, Substrate specificity of the DNA unwinding activity of the RecBC enzyme of Escherichia coli. J. Mol. Biol., 185, 431–443 (1985).

36. M.C. Dillingham, S.C. Kowalczykowski, RecBCD enzyme and the repair of double-stranded DNA breaks. Microbiol. Mol. Biol. Rev., 72, 642–71 (2008).

37. C. Masterson, P.E. Boehmer, F. McDonald, S. Chaudhuri, I.D. Reconstitution of the activities of the RecBCD holoenzyme of Escherichia coli from the purified subunits. J. Biol. Chem., 267, 13564–13572 (1992).

38. M. Yu, J. Souaya, D.A. Julin, The 30-kDa C-terminal domain of the RecB protein is critical for the nuclease activity, but not the helicase activity, of the RecBCD enzyme from Escherichia coli. Proc. Natl. Acad. Sci. USA, 95, 981–986 (1992).

39. J.-Z. Sun, D.A. Julin, and J.-S. Hu, The nuclease domain of the Escherichia coli RecBCD enzyme catalyzes degradation of linear and circular single-stranded and double-stranded DNA. Biochemistry, 45, 131–140 (2006).

40. J. Wang, R. Chen, D.A. Julin, A single nuclease active site of the Escherichia coli RecBCD enzyme catalyzes single-stranded DNA degradation in both directions. J. Biol. Chem., 275, 507–513 (2000).

41. M.R. Singleton, M.S. Dillingham, M. Gaudier, S.C. Kowalczykowski, D.B. Wigley, Crystal structure of RecBCD enzyme reveals a machine for processing DNA breaks. Nature, 432, 187–193 (2004).

42. N. Garcia-Quintans, L. Bowden, J. Berenguer, M. Mencia, DNA interference by mesophilic argonaute CbcAgo. F1000 Research, 8, e321 (2020).

43. L. Greenough, K.M. Schermerhorn, L. Mazzola, J. Bybee, D. Rivizzigno et al. Adapting capillary gel electrophoresis as a sensitive, high-throughput method to accelerate characterization of nucleic acid metabolic enzymes. Nucl.Acids Res., 44, e15 (2016).

44. J. Gerritsen, S. Fuentes, W. Grievink, L. van Niftrik, B.J. Tindall et al. Characterization of Romboutsia ilealis gen. nov., sp. nov., isolated from the gastro-intestinal tract of a rat, and proposal for the reclassification of five closely related members of the genus Clostridium into the genera Romboutsia gen. nov., Intestinibacter gen. nov., Terrisporobacter gen. nov. and Asaccharospora gen. nov. J. Syst. Evol. Microbiol., 64, 1600–1616 (2014).

45. K. Umezu, and H. Nakayama, RecQ DNA helicase of Escherichia coli. Characterization of the helix-unwinding activity with emphasis on the effect of single-stranded DNA-binding protein., J. Mol. Biol., 230, 1145–1150 (1993).

46. B. Rad, A.L. Forget, R.J. Baskin, S.C. Kowalczykowski, Single-molecule visualization of RecQ helicase reveals DNA melting, nucleation, and assembly are required for processive DNA unwinding. Proc. Natl. Acad. Sci. USA, 112, E6852–E6861 (2015).

47. E.A. Hunt, E. Tamanaha, K. Bonanno, E.J. Cantor, N.A. Tanner, Profiling Thermus thermophilus Argonaute guide DNA sequence preference by functional screening. Front.Mol.Biosci., 8, 670940 (2021

48. S. Willkomm, C.A. Oellig, A. Zander, T. Restle, R. Keegan, Structural and mechanistic insights into an archaeal DNA-guided Argonaute protein. Nat. Microbiol., 2, 17035 (2017).

49. Y. Wang, G. Sheng, S. Juranek, T. Tuschl, D.J. Patel, Structure of the guide-strand-containing argonaute silencing complex. Nature, 456, 209–213 (2008).

50. R.C. Edgar, MUSCLE: multiple sequence alignment with high accuracy and high throughput. Nucleic Acids Res., 32, 1792–1797 (2004).

51. S.R. Eddy, Accelerated profile HMM searches. PLOS Comp. Biol., 7, e1002195 (2011).

## Reference

1. K.D. Raney, L.C. Sowers, D.P. Nillar, S.J. Benkovic, A fluorescence-based assay for monitoring helicase activity. Proc. Natl. Acad. Sci. USA, 91, 6644–6648 (1994).

